# Long-term surveillance reveals hybridization by nuclear reassortment and intercontinental spread as major evolutionary drivers in wheat yellow rust

**DOI:** 10.1101/2025.05.22.655633

**Authors:** Mogens Støvring Hovmøller, Tine Thach, Julian Rodriguez-Algaba, Jens Grønbech Hansen, Marcel Meyer, David Hodson, Kumarse Nazari, Robert F. Park, Rita Tam, Mareike Möller, Benjamin Schwessinger, John P. Rathjen, Paula Silva, Venancio Riella, Annemarie Fejer Justesen

## Abstract

- Evolutionary forces affecting crop pathogens, including hybridization and long-distance dispersal (LDD), may have strong implications for food security and sustainable plant disease control at global scales. However, consolidated evidence is often lacking due to absence of consistent pathogen surveys beyond national capacities.
- Our study documents world-wide connectivity between populations of *Puccinia striiformis*, causing yellow rust on cereals and grasses, when analysing 3240 pathogen samples collected in 41 countries on six continents from 2009-2023. Our analyses revealed twelve cases of inter-continental spread of *Puccinia striiformis*, including seven cases with major impact on disease epidemics in recipient areas.
- Somatic hybridization by nuclear reassortment between co-existing multi-locus genotypes (MLGs) on a common host was the most plausible mechanism for the emergence of three novel clonal groups of global relevance, first detected in Europe. Subsequently, onwards spread to South America and Australia was observed. Several high-impact incursions from South Asia into East Africa were also observed, including a genotype with a dramatic impact on wheat breeding programs of global relevance.
- Our study stresses an urgent need for coordinated crop pathogens monitoring across borders. Only global efforts will enable prevention and control of pathogens that represent major challenges for food security at regional and global scales.

## Introduction

An increase in human travel and trade and the escalating speed of changing climate and weather patterns have renewed attention on disease epidemiology in humans, animals and crop plants, e.g., by the US National Academies of Sciences (Olsen et al., 2011), and the United Nations Food and Agriculture Organisation (FAO, 2024). Long distance dispersal (LDD) is an important epidemiological feature of many pathogens, including those affecting crops, because even rare LDD events can have a major impact on crop production (Brown and Hovmøller, 2002; Carvajal-Yepes et al., 2019). Recent reviews suggest that LDD events of cereal rust pathogens are increasing in frequency at the intercontinental scale, possibly due to intensified crop production, climate change resulting in milder winters in temperate regions, and increased travel and commerce (Park, 2015; Jin et al., 2020; Hovmøller et al., 2023). More frequent LDD events may delay or prevent achievement of the 70% increase in food and agricultural production required by 2050 to meet global population growth (van Dijk et al., 2021).

One major obstacle for the detection of LDD events in plant pathology is the lack of regular and extensive pathogen population surveys beyond national capacities (Jeger and Pautasso, 2008; Ristaino et al., 2021), including common sampling protocols, harmonised population genetic data and analytic tools (Patpour et al., 2022). This makes it difficult to establish the exact geographical and evolutionary origins of a new pathogen variant as it spreads into recipient areas. Multiple tools are available for analyses of population genetic features such as genetic distance, population subdivision and migration rates within and among pathogen populations (Zhan et al., 2003; Ali et al., 2014a; Bebber et al., 2014). In predominantly clonal populations with a high degree of preserved haplotypes, the tracking of individual multi-locus genotypes (MLGs) may represent a different and powerful strategy for the detection of LDD events (Brown and Hovmøller, 2002; Hovmøller et al., 2002).

The rust fungi (*Pucciniales*) are the most species rich and complex order of plant pathogens, consisting of almost 8000 species. They are characterized by a true biotrophic lifestyle and are individually adapted to their host plant at species and cultivar levels (Flor, 1971; Anikster, 1985; Aime et al., 2017). *Puccinia striiformis* f. sp. *tritici (Pst)*, causing yellow (or stripe) rust on wheat, is a dikaryotic, heteroecious and macrocyclic fungus requiring a primary cereal/grass host and a taxonomically unrelated alternate host, e.g., *Berberis* spp., to complete its life cycle (Jin et al., 2010).

Multiple studies have reported high diversity in *Pst* populations in the near-Himalayan region (including China, Pakistan, Nepal), which is considered the center of diversity of this fungus, e.g., (Duan et al., 2010; Ali et al., 2014b; Thach et al., 2016; Huang et al., 2021), however, the functional role of the sexual host, *Berberis spp.,* in these areas is not fully understood because *Pst* is rarely observed on *Berberis spp.* in nature (Jin et al., 2010; Berlin et al., 2013; Wang et al., 2016; Rodriguez-Algaba et al., 2022).

In contrast, many studies in other regions have documented a clonal *Pst* population structure, where spontaneous and high mutation rates coupled with host-induced selection are major evolutionary driving forces, e.g., in Europe (Hovmøller et al., 2002; Enjalbert et al., 2005; Hovmøller and Justesen, 2007; de Vallavieille-Pope et al., 2012b; Hovmøller et al., 2016; Saunders et al., 2019), Australia (Wellings and McIntosh, 1990; Steele et al., 2001; Ding et al., 2021), South America (Anibal Carmona et al., 2019), North America (Markell and Milus, 2008; Brar et al., 2018), as well as Central- and West Asia and North- and East Africa (Hovmøller et al., 2008; Ali et al., 2014a; Thach et al., 2016; Walter et al., 2016; Ali et al., 2017b). Several studies involved comparative analyses between high-diversity areas in the near-Himalayan region, including China, and clonal populations from elsewhere, revealing clear geographic population structures with limited overlap. At the same time, rare LDD events have ensured some degree of connectivity across large time and spatial scales, e.g., Hovmøller et al. (2008); (Ali et al., 2014a; Anibal Carmona et al., 2019; Ding et al., 2021; Župunski et al., 2024). In 2011, clonal populations in Europe were influenced by incursions likely originating from populations in the near-Himalayan region, including a number of distinct MLGs with very different virulence- and molecular patterns compared with the pre-existing population that were largely replaced within few years (Hovmøller et al., 2016).

So far, somatic hybridization by fusion of hyphae, which may lead to nuclear reassortment where parental haplotypes are preserved, has only been reported under experimental conditions in *Pst* (Little and Manners, 1967; Little and Manners, 1969; Lei et al., 2017), unlike for the wheat leaf rust pathogen *P. triticina* (Park and Wellings, 2012; Sperschneider et al., 2023), the wheat stem rust pathogen *P. graminis* f. sp. *tritici* (Li et al., 2019), and *P. coronata* causing oat crown rust (Henningsen et al., 2024).

The increased focus on *Pst* epidemiology at the global level since 2000 coincided with the emergence of *P. graminis* f. sp. *tritici* (*Pgt*) race “Ug99” in East Africa that was virulent on more than 80% of wheat lines developed by The International Maize and Wheat Improvement Center (CIMMYT) (McIntosh and Pretorius, 2011). In response, the Borlaug Global Rust Initiative (BGRI) was formed and laid the foundation for more extensive and regular wheat rust disease and pathogen surveys. These included Africa and Asia, which were connected with national rust survey programs in Europe and elsewhere by the establishment of the Global Rust Reference Center in 2008 (Hovmøller, 2011). This resulted in commonly agreed sampling and genotyping methodologies for these two wheat rusts, i.e., often using a core set of simple sequence repeat (SSR) or single nucleotide polymorphic (SNP) markers (Bahri et al., 2008; Patpour et al., 2022; Szabo et al., 2022; Thach et al., 2025), and more recently MARPLE (Radhakrishnan et al., 2019; Župunski et al., 2024), which generally have allowed a high degree of alignment of results from studies representing different time periods and sampling areas.

The combined international efforts have allowed analyses of the spread and evolution of *Pst* and *Pgt* across borders and continents, including the return of *Pgt* into Europe and connectivity to epidemics elsewhere (Patpour et al., 2022), including Western Siberia 2015-2016 (Shamanin et al., 2016) and East Africa (Olivera et al., 2015). In addition, independent surveys have reported large scale epidemiological consequences of new incursions of *Pst* into South America in 2017 (Anibal Carmona et al., 2019; Riella et al., 2024) and Australia in 2017-2018 (Ding et al., 2021).

The present study investigated drivers of *P. striiformis* evolution at global scale based on genotypic analyses of 3240 samples; 3180 samples collected at four continents (37 countries) between 2009 and 2023, complemented by genotype information of additional 60 samples representing long-term surveillance efforts in Europe, North America (Milus et al., 2015a) and Australia (Ding et al., 2021). LDD events were identified by investigating the potential sharing of genotypes in putative source and recipient areas, considering the time of first detection as well as accumulation of genetic diversity in the considered populations. In addition, the plausibility of wind dispersal along hypothesized transmission routes was explored by conducting trajectory simulations considering typical spore survival times and mean wind patterns. Likely evolutionary origin of successful LDD migrants were explored in detail in specific cases, including hypothetical hybridization events by nuclear reassortment, which have not previously been demonstrated in *Pst* under non-experimental conditions. The prospects for prevention and mitigation efforts to reduce the increasing impact of crop pathogen spread at intercontinental scale are discussed.

## Materials and Methods

### Collection and recovery of yellow rust samples

Pathogen sampling reflected the coordinated efforts in wheat rust surveillance facilitated by research programs within the Borlaug Global Rust Initiative and European Union research and network initiatives, collectively defining seven geographic *Pst* population between 2009-2023, as well as reference populations in North America and Australia, the latter representing the main clonal divergences in *Pst* since first incursion into Australia in 1978 **(Fig. S1).**

A total of 3180 time- and geo-referenced samples of yellow rust mainly from wheat (and occasionally other cereals and grasses) were collected in 37 countries on four continents between 2009 and 2023 (Table S1). The number of samples varied across years and locations, reflecting prevalence of epidemic outbreaks and available sampling efforts. The Regional Cereal Rust Research Center (RCRRC), ICARDA, Turkey provided 213 samples from the Middle East and West Asia. An additional pre-existing 60 isolates served as references of previously identified *Pst* groups, and *Pst* populations in USA (Milus et al. (2015b) and Australia (Ding et al. (2021), i.e., the study comprised 3240 samples in total.

Recovery of incoming samples and multiplication of urediniospores followed the procedure by Thach et al. (2025), after which they were harvested, dried, and stored in liquid nitrogen (−196°C) or freezer (−80°C) until further use. .

### DNA extraction and SSR genotyping

From 2009 to 2012, DNA was extracted from dried urediniospores (approx. 20 mg/sample) according to Thach et al. (2016). Since 2013, extraction was based on dried leaf segments with single rust lesions taken from multiplication plants or directly from incoming samples according to Thach et al. (2025). Genotyping was done using 19 simple sequence repeat (SSR) co-dominant markers and allele sizes were manually scored, allowing up to two missing markers per sample (Thach et al., 2025). Samples with allele sizes and/or combinations suggesting more than a single genotype were excluded from the dataset.

Detection of novel or exotic genotypes in a region/country was confirmed by additional independent DNA extractions from original sample material and genotyping. Two hundred thirteen samples from the Middle East (2018-2020) collected by Turkey-ICARDA Regional Cereal Rust Research Center were SSR genotyped by EUROFINS, France. Scoring of allele sizes and data alignment to existing format were carried out at the Global Rust Reference Center (GRRC). Assignment of multilocus genotype identity (MLG #) was done for samples without missing data using poppr package version 2.9.3 implemented in the R environment (Kamvar et al., 2014; Kamvar et al., 2015); for samples with missing data at a maximum of two marker loci, new MLGs were considered in case scored allele sizes revealed additional variability, whereas missing observations were not taken into account for MLG identity. Duplicate MLGs derived from the same incoming sample were removed from the dataset.

### Clustering within clonal groups

The clustering within 19 previously defined clonal Pst groups (PstS0-PstS18), each consisting of closely related MLGs and associated races (Thach et al., 2025), were used to illustrate population differences at geographical scale and time. The definition and naming of new clonal groups have been inspired by the potential epidemic impact across large areas (Hovmøller et al., 2010), significant divergence from previous clonal groups and/or group-specific diagnostic markers (Walter et al., 2016; Ali et al., 2017b). An updated summary of named *Pst* groups is available at the GRRC website: https://agro.au.dk/forskning/internationale-platforme/wheatrust/yellow-rust-tools-maps-and-charts/definitions-of-races-and-genetic-groups.

### Population diversity and phylogenetic analysis

The discriminative power of applied markers was illustrated by a genotype accumulation curve calculated in the poppr R package based on samples without missing data (*n*= 2588), and random resampling 10,000 times without replacement (Kamvar et al., 2014) (Fig. S2).

The dataset for phylogenetic analysis was compiled by including one representative sample from each MLG (clone corrected). An unrooted phylogenetic Neighbor Joining tree was constructed to visualize global diversity within *Pst*, including the positions of major clonal groups (PstS0-PstS18). The calculations were based on genetic distances based on the poppr package version 2.9.3 implemented in the R environment, 1200 bootstraps and cutoff at 50 (Kamvar et al., 2014), where Bruvo’s distance for stepwise mutation for microsatellites was calculated according to Bruvo et al. (2004).

### Whole genome confirmation of a somatic hybridization event

The hypothesized somatic hybridization resulting in PstS10 inferred by SSR and PstS0 genome data was validated by two newly generated reference assemblies of isolates representing PstS7 and PstS10, respectively. The fully phased nuclear-assigned reference genome of an Australian sampled PstS10 isolate (Ding et al. 2021) was generated as described previously for PstS0 (Tam et al., 2025) with the exception of using PacBio HiFi and Oxford Nanopore Technology (ONT) R10.4 based long-reads and Hi-C as input for Verkko genome assembler (Rautiainen et al., 2023). The haplotype-phased reference genome of PstS7 used ONT R10.4.1 simplex reads and Hi-C with hifiasm --ont (Cheng et al., 2025). To expand upon the MLG results at the genome level, comparative alignments were performed between the nuclear-phased haplotypes of a proposed somatic hybrid, PstS10, and those of its two hypothetical parental isolates, PstS0 (Tam *et al*., 2025) and PstS7. Pairwise whole-genome alignments were conducted using NUCmer v3.1 as part of the MUMmer package (Marçais et al., 2018) with default settings. Only alignment blocks with >75% identity and longer than 500 bp were retained. SNP counts and average identity were computed from 1-to-1 alignments using MUMmer’s DNAdiff v1.3. Dot plots were generated using a script adapted from paf2dotplot.r (https://github.com/moold/paf2dotplot). The chromosomal locations and allele sizes for PstS7 and PstS10 are visualized in Table S2 and Fig. 3, respectively, and calculated amplicon lengths were compared with SSR genotyping results.

### Tests for somatic hybridization by nuclear reassortment

The hypothesis of evolution of three novel MLGs by somatic hybridization and nuclear reassortment among corresponding parental MLGs was explored according to a series of connected steps (Fig. S3), taking advantage of haplotype-phased genome data of a PstS0 isolate (Plant Breeding Institute accession number 415 collected in 1982) (Schwessinger et al., 2018; Tam et al., 2025) followed by the generation of nuclear-phased genomic data of a PstS7 and a PstS10 isolate (this study). Nuclear-specific SSR haplotypes of hypothetical parental and hybrid isolates were determined by blast analysis (-word_size 12 -evalue 1 - dust no -outfmt 6) of 20 primer sequence pairs against the PstS0 genome (Table S2) and visualized according to Wolfe et al. (2013). The calculated SSR haplotypes were adjusted to observed (scored) allele sizes to allow inference of observed parental and hybrid haplotypes of PstS13, PstS14, and PstS18, respectively, the latter reported recently as a likely hybrid where only one parent has been identified, following the approach and procedures according to Fig. S3.

### Virulence phenotyping

Recovered, hypothesized migrant isolates from putative source and recipient areas were virulence phenotyped to provide an independent assay for genetic relatedness. An updated protocol using a 0-9 disease scoring scale according to (McNeal et al., 1971), where infection types (IT) from 0-6 were generally considered incompatible (avirulent) and IT 7-9 compatible (virulent) was followed (Thach et al., 2025). The interpretation of virulence/avirulence was generally based on two (or more) independent wheat lines carrying the considered *P. striiformis Yr* resistance gene.

### Plausibility of wind-dispersal

Plausibility of wind-dispersal was assessed based on (i) reviewing data from previous studies on related fungal pathogens and other microorganisms (Brown and Hovmøller, 2002; Isard et al., 2005; Isard et al., 2011; Meyer et al., 2017a; Meyer et al., 2017b; Prank et al., 2019; Bradshaw et al., 2022; Radici et al., 2022; Brodsky et al., 2023; Miedaner and Garbelotto, 2024); (ii) analyzing long-term mean wind patterns (wind direction and speed at times of overlapping wheat seasons at source and recipient areas; International Research Institute for Climate and Society (https://iridl.ldeo.columbia.edu/maproom/Global/Climatologies/Vector_Winds.html); (iii)estimating typical pathogen survival times from literature (Maddison and Manners, 1972; Rapilly, 1979; Aylor, 1986); and (iv) conducting a set of atmospheric trajectory simulations using HYSPLIT (Stein et al., 2015) with two different global-scale gridded meteorological datasets [NOAA’s GDAS, 3-hourly temporal resolution and 1 degree horizontal spatial resolution; ECMWF’s ERA5, hourly temporal resolution and 0.25 degree spatial resolution (Hersbach et al., 2020). Time-backwards trajectory simulations were conducted from locations in recipient areas with GDAS meteorological input data to investigate origins of air masses that could have transported spores to recipient areas where they were first detected. Time-forward simulations were conducted with ERA5 meteorological input to analyze the plausibility of stepwise airborne transmission along two selected pathways (Europe to South America via Northern Africa; and Central Asia to East Africa via the Middle East).

## Results

### Genotypic *Pst* diversity at global scales

A majority of the samples investigated (91.7%) grouped into previously defined clonal groups PstS0-PstS18 (Fig. 1). The remaining samples consisted of 74 diverse MLGs mainly from South Asia, where high diversity has been reported previously, and from non-wheat hosts in northern Europe, e.g., barley, wild grasses, and triticale. The 20 reference samples from North America contained six MLGs within PstS0, PstS1_2, PstS18, and two unnamed groups, respectively. The 31 reference samples from Australia were embedded in seven MLGs that grouped within PstS0, PstS1_2, PstS10 and PstS13, respectively, whereas six reference samples from Europe and one from East Africa (1960-2006) represented PstS1_2, PstS3 and PstS4.

**Fig. 1.**
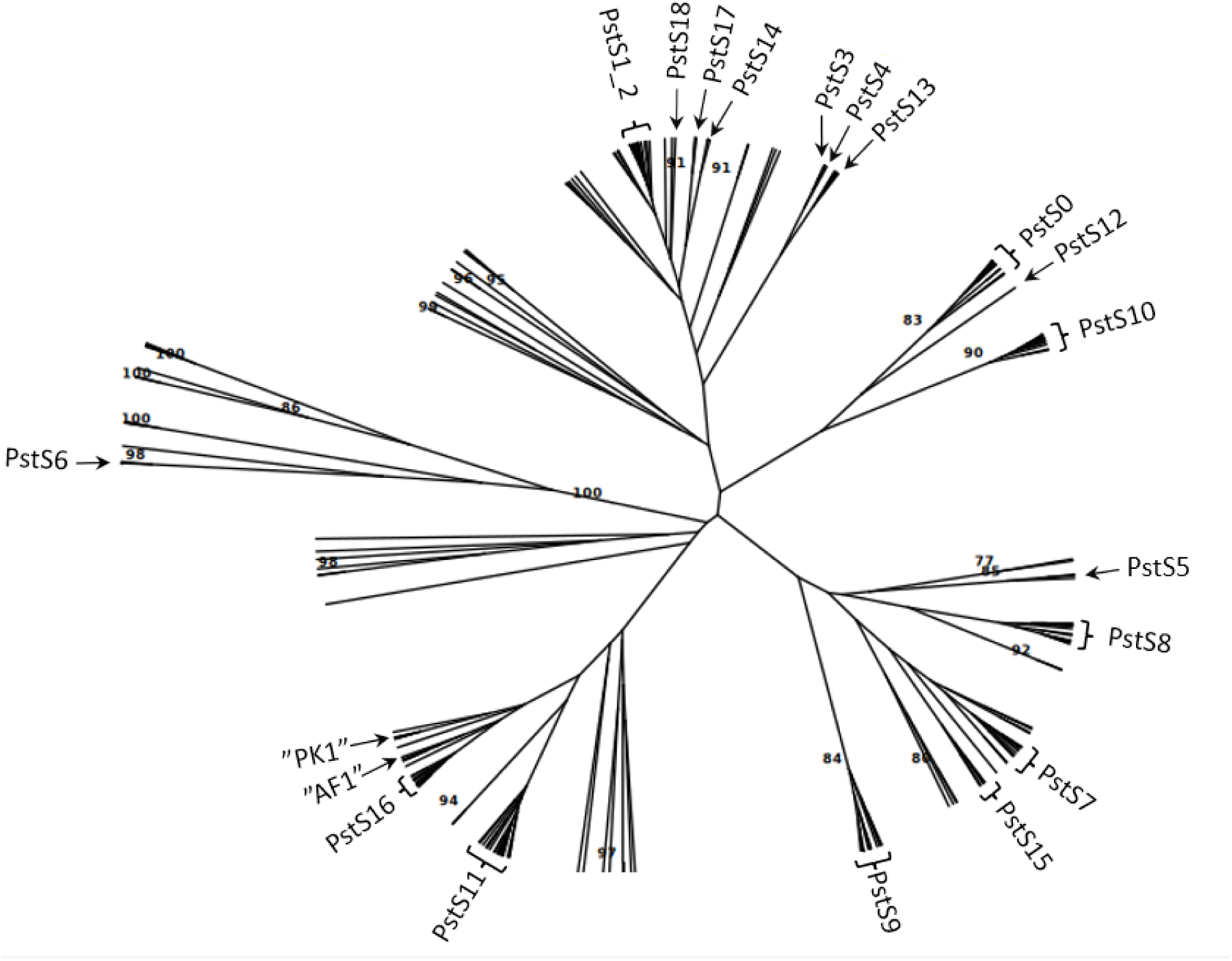
Unrooted Neighbor Joining tree (clone-corrected) using Bruvo’s distance showing the genetic relationship of 244 multi-locus genotypes (MLGs) representing world-wide sampling of *P. striiformis* in 41 countries on six continents (2009-2023), including reference isolates representing populations in Australia (1978-2021) and North America (1972-2023). Position of named clonal groups (PstS0-PstS18) and two temporary named groups (AF1, PK1), consisting of clusters of closely related MLGs, are indicated. Remaining branches mainly represent MLGs from diverse populations in South Asia and Central/West Asia.

Diversity within clonal groups varied considerably, reflecting time since first detection of founder MLG (evolutionary time) as well as disease prevalence and related sampling intensity (Table S1; Fig. S1). The highest number of MLGs were observed in PstS1_2, first detected in East Africa in the 1970s. Yet the highest diversities relative to number of samples were observed for PstS7, PstS8 and PstS9 when considering clonal groups, defined according to Thach et al. (2025), with more than 25 samples (Table S2). Unique MLG identification numbers with reference to hybridization and LDD events shown in right column.

The prevalence of named clonal groups (PstS0-PstS18) changed significantly over time and across geographical areas (Northern Europe, Middle East, East Africa, Northwest Africa, South Asia, Central-West Asia, South America and Australia), suggesting geographic subdivisions and dynamic pathogen populations during time (Fig. 2).

**Fig. 2.**
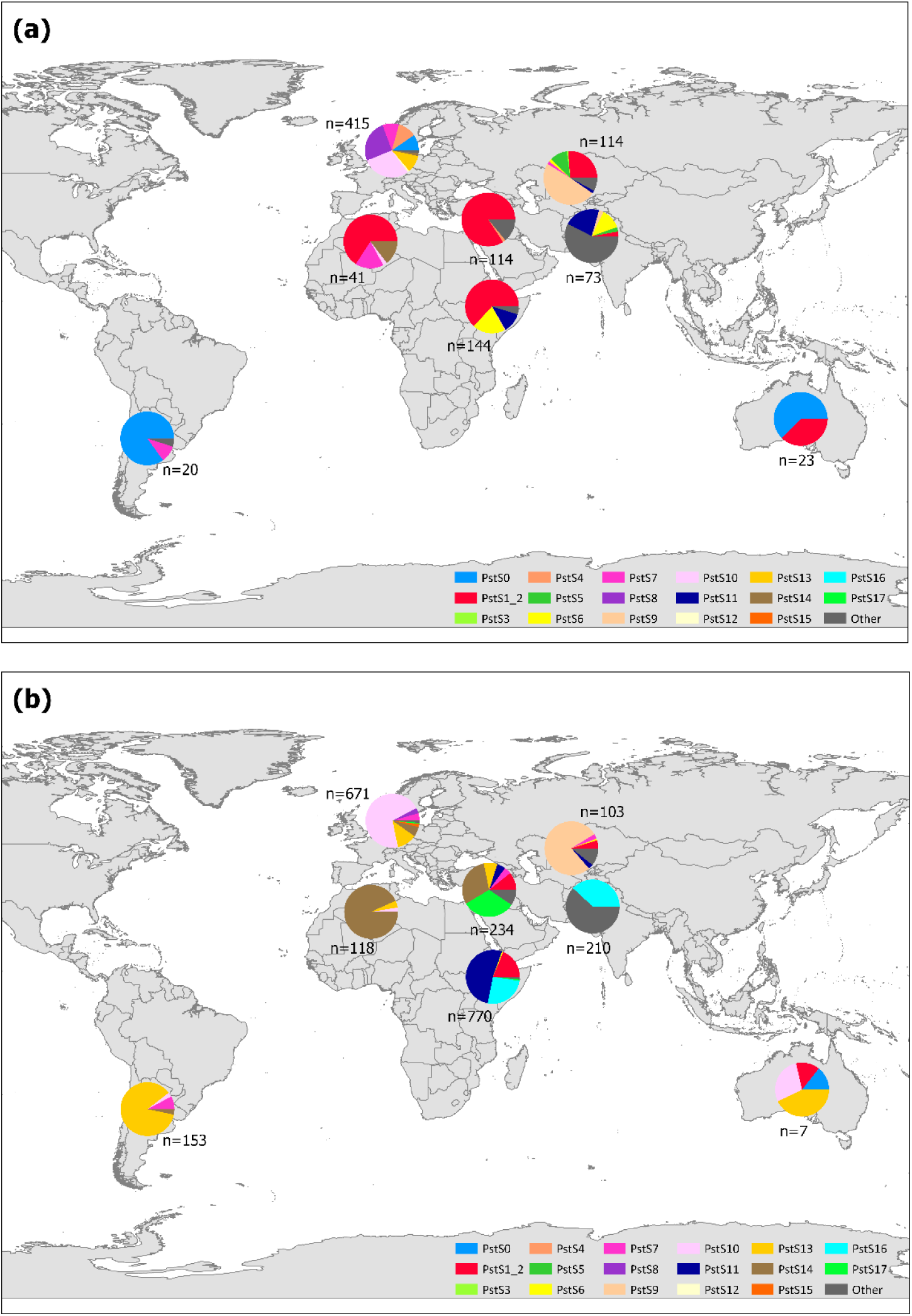
Frequency of *P. striiformis* clonal groups in distinct geographical populations covering major wheat growing regions world-wide between 2009 and 2023 (cf. Table S1). A) Sampling period 2009-2016, and B) sampling period 2017-2023, (*n*= sample size). Australian data represent key reference samples 1978-2021, i.e., from first *Pst* disease appearance on the continent and until recent time. North American reference data not shown due to low sample size and uneven representation over time and area.

The diverging geographical distribution of clonal groups was striking, e.g., PstS1_2 (high temperature-adapted) was prevalent across Africa and the Middle East, but absent from South America, Europe and South Asia. PstS5 and PstS9 were only detected in Central Asia, and PstS0 was only detected in Europe, the Americas and Australia. PstS8 was prevalent in northern Europe until 2016 but has not detected in other areas. PstS10 was the most prevalent group in Europe from 2014 and onwards but was not detected in East Africa and South- and Central Asia.

The dataset revealed several cases of emergence of novel clonal groups with significant epidemiological impact on global scale. For example, the PstS10 founder MLG was first detected in Europe in 2012 and became widespread, accounting for a total of 82% of PstS10 samples in the study (Table S2). This founder MLG was followed by the emergence of a further 14 closely related MLGs, consistent with clonal evolution by mutation. The founder MLG of PstS13 (# 21) was only detected in Europe during 2015-2016, and was first observed in South America in 2017, where yellow rust re-appeared at epidemic levels after many years without significant impact. The emergence of a total of six closely related MLGs in PstS13 revealed another case of typical clonal evolution. The likely founder MLG of PstS14 (#138, 72% of samples, Table S2), which was first detected in NW Africa in 2016, represented a third novel clonal group in the dataset.

### Exploring the hypothesis of somatic hybridization

The founder MLGs of these three novel clonal groups diverged significantly from previous clonal groups and MLGs, and were not resampled outside Europe until 2-3 years after first detection. We therefore explored the hypothesis of hybridization of co-existing parental isolates on common host plants by nuclear reassortment according to Fig. S3. An initial screen of isolates of named clonal groups that could potentially contribute to a somatic hybridization resulting in the founder MLG of PstS10 (#244, Table S2) revealed that all except PstS0 and PstS7 isolates were excluded due to mismatching allele sizes (Table S3).

To support our initial observation of PstS0 and PstS7 as potential nuclear donors for PstS10, a comparative analysis of the chromosomal locations of the 19 SSR markers was performed. This analysis made use of existing and newly generated haplotype phased reference genome assemblies of PstS0 (Tam et al. 2025) and PstS7 and PstS10 (this study, Table S4, Fig. 3), respectively.

**Fig 3.**
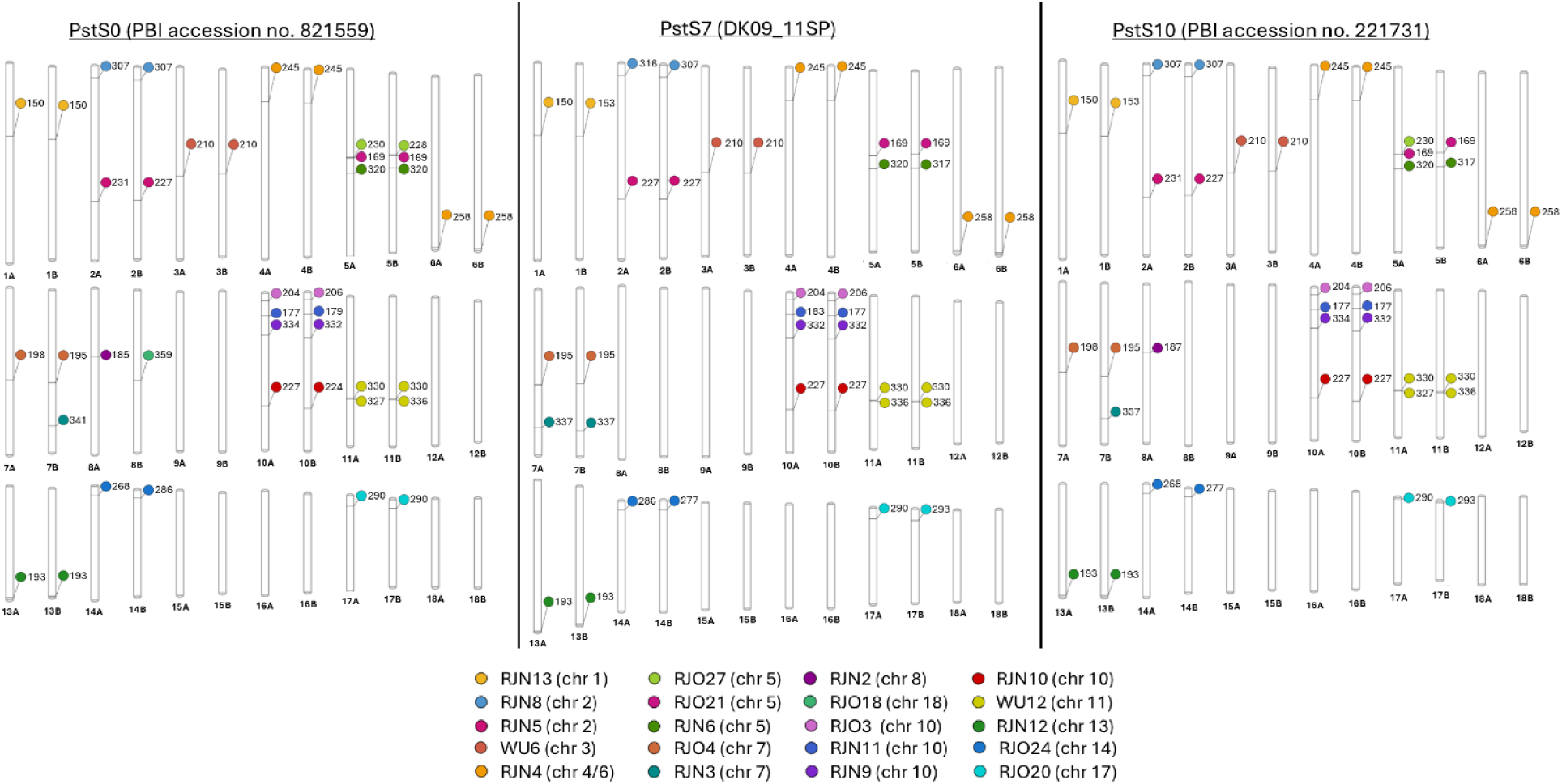
SSR marker placement on haplotype-phased genomes of PstS0, PstS7 and PstS10 determined by Blast analyses using pairwise SSR primer sequences. Markers were distributed across 13 chromosomes and detected in both nuclei (denoted A and B) except for RJN2 and RJO27. RJO3 (chromosome 10) was generally monomorphic and not scored in this study.

Interestingly, the SSR markers were distributed on 13 of 18 assigned *Pst* chromosomes, generally detected in both nuclei, except for RJN2, RJN3 and NJO27, resulting in a set of well-documented markers with a wide representation of the *Pst* genome. The SSR haplotypes derived from genome data, and the corresponding haplotypes deduced from the observed MLGs in the dataset, resulted in a full match based on a historic prevalent MLG #277 of PstS0 in Europe, MLG #321 in PstS7, first detected in Europe in 2011, and the founder MLG of the resulting hybrid in PstS10, i.e., #244 (Table S2).

The SSR haplotype results were confirmed by pairwise genome alignments and SNP counts derived from the nuclear-phased haplotypes of the three isolates representing PstS0, PstS7 and the founder MLG of PstS10, respectively (Fig. 4). Nucleus A in the hybrid (PstS10) displayed a 99.98% similarity with the corresponding nucleus A in PstS0 based on whole genome alignments with only ∼3k SNPs differences when compared to intra isolate nuclear similarities of less than 98.35% with over 650k SNPs in PstS0. Similarly, the nucleus B of the hybrid displayed 99.98% similarity with the corresponding nucleus in PstS7 with about ∼ 1k SNPs when compared to over 150k SNPs between the two nuclei within PstS7.

**Fig. 4.**
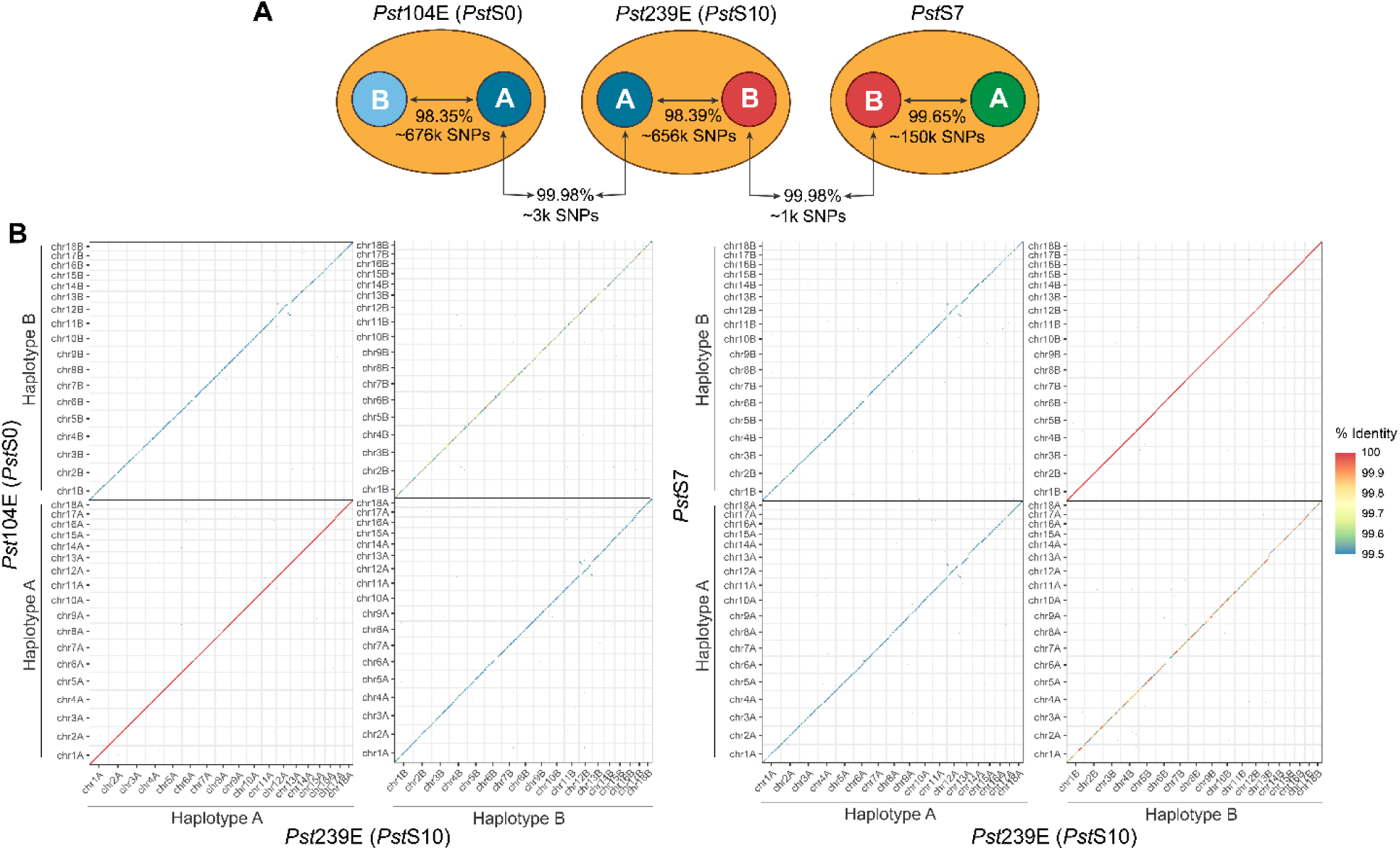
A. Total SNP counts and average identity from pairwise genome alignments among the nuclear-phased haplotypes of a proposed somatic hybrid (PstS10), and two hypothetical parental isolates from PstS0 and PstS7, respectively. B. Dotplots of all pairwise genome alignments among the nuclear haplotypes. Only alignment blocks with >99.5% identity and longer than 10 kbp are shown.

An overlay of allele sizes of the PstS7 haplotypes on PstS13 and PstS14 suggested that the PstS7 nucleus B was likely involved in additional hybridization events that resulted in the founder MLGs of these (Table S2). Rationale and conclusions about specific SSR haplotypes, MLG variants and likely 2^nd^ parents are provided in the discussion, where co-existence of hypothetical parental variants in time and space is taken into account.

### Long distance dispersal events

Twelve cases of *Pst* incursions at intercontinental scales (I-XII) were documented by shared MLGs and shared virulence phenotype in case alive isolates were available for testing (Table 1, Fig. 5). PstS13 that was first detected in Europe in 2015 (MLG #21) spreads into South America in 2017 and Australia in 2018 (Table 1, dispersal events I and III). Recovered samples from South America (2017) shared virulence phenotype with a representative sample of the founder MLG (#21) in Europe. In 2017, PstS14 was detected for the first time outside Europe (South America, dispersal events II), and PstS10 was first detected in Australia (dispersal event IV).

**Table 1.**
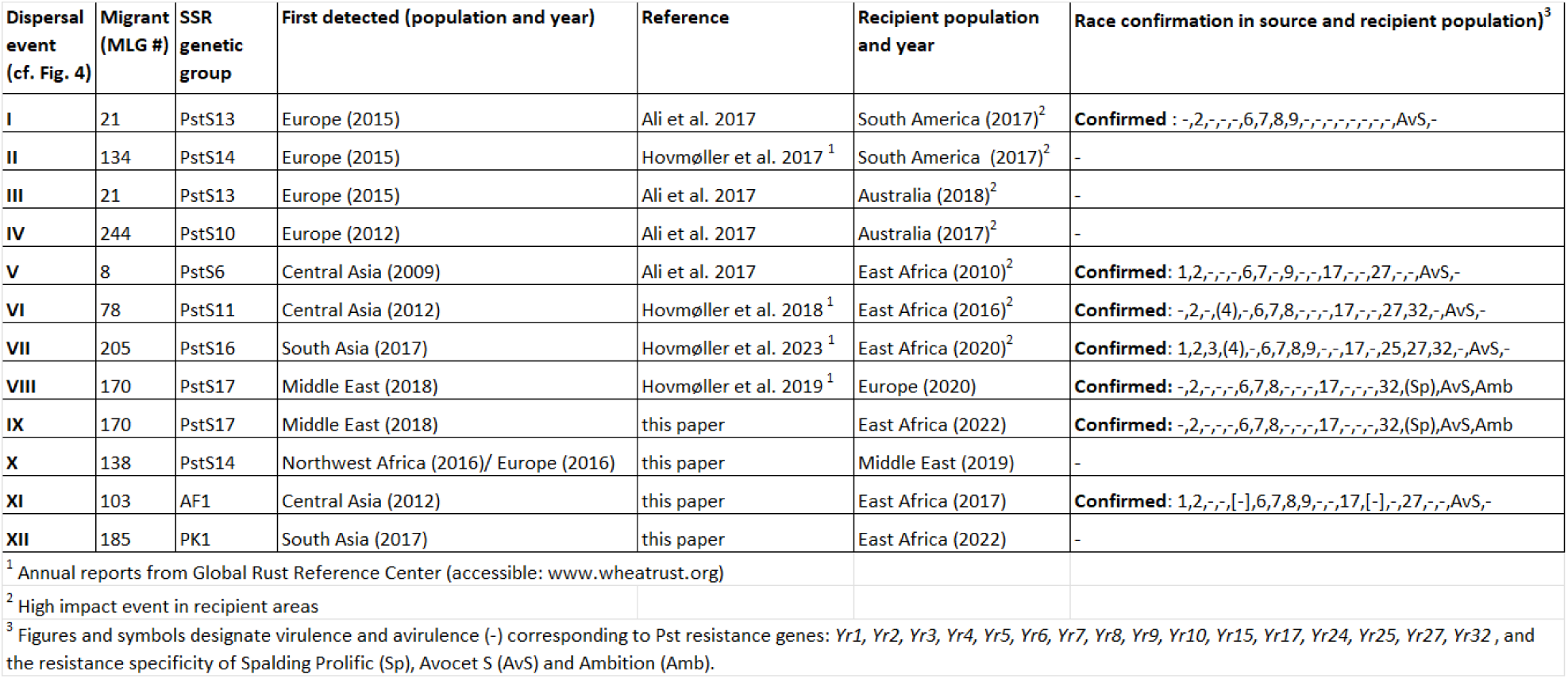
Intercontinental *Puccinia striiformis* dispersal events (I-XII) within a 15-year period (2009-2023) documented by source and recipient areas for clonal groups, unique multi-locus genotypes (MLGs) within these, and race phenotype in case alive isolates were available for testing.

**Fig. 5.**
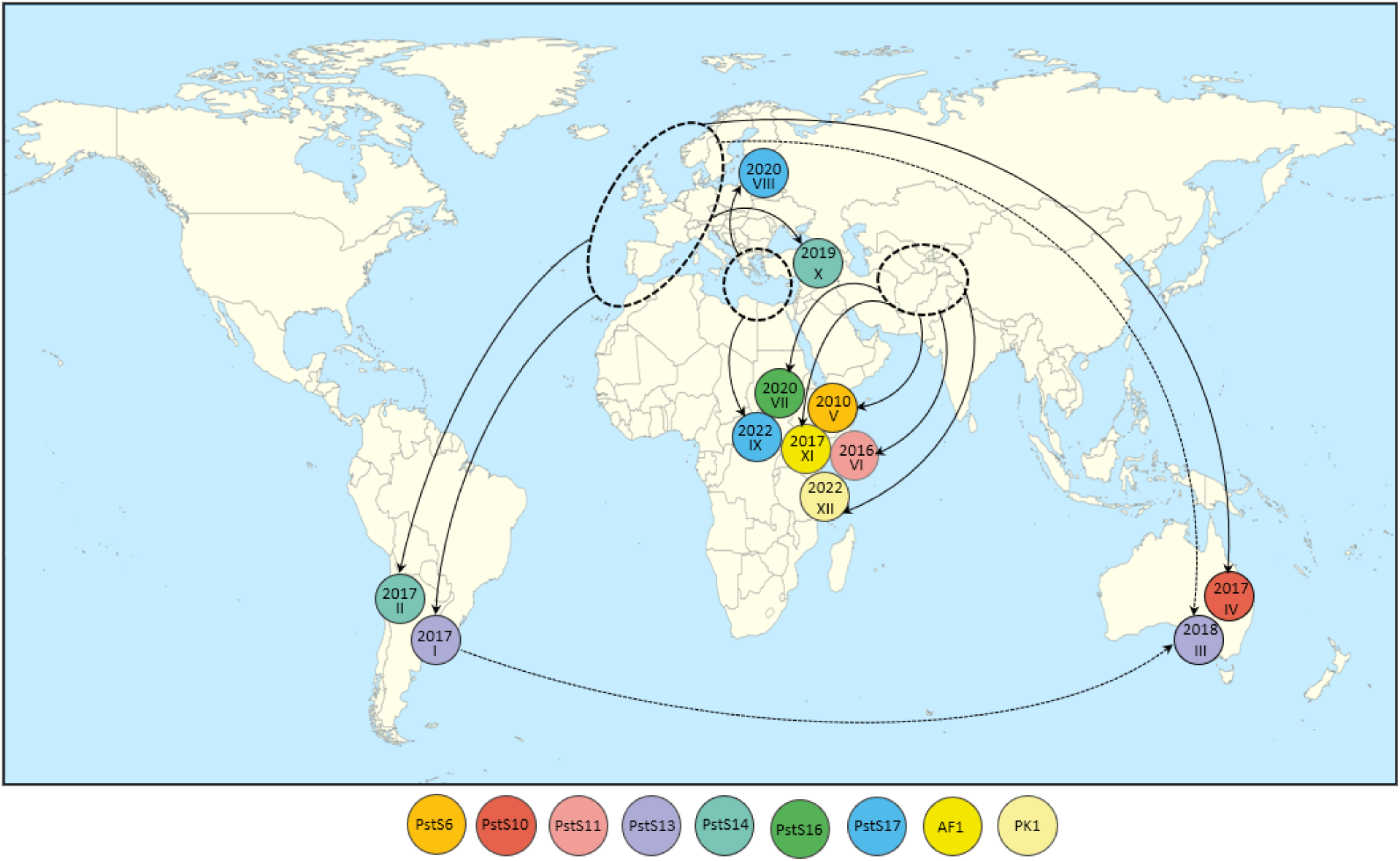
Documented intercontinental incursions of *P. striiformis* 2009-2023, based on sequential resampling of MLGs/pathogen races on different continents. Dotted circles represent the putative source areas where considered *Pst* clonal groups and MLGs defined by SSR markers were first detected, typically 2-3 years ahead of first detection in recipient areas. Putative transmission routes indicated by black arrows, colored circles represent first year of detection on recipient continent and corresponding dispersal events. Dotted arrows: PstS13-MLG #21 was present at two continents at time of incursion into Australia. The positions of circles and arrows on the map are indicative.

PstS6 (MLG #8), PstS11 (MLG #78) and PstS16 (MLG #205) that were first detected in Central/South Asia in 2009, 2012 and 2017, respectively, were observed for the first time in East Africa in 2010, 2015 and 2020, respectively (events V, VI, VII, Table 1). PstS17 (MLG #170), first detected in the Middle East in 2018, was detected in northern Europe in 2020 (event VIII, Table 1) and East Africa in 2022 (event IX). Identical virulence phenotype based on pairs of recovered samples representing source and recipient areas for dispersal events V – IX confirmed a shared, common origin in each of these cases. PstS14 (MLG #138) was first detected in the Middle East in 2019 (X, Table 1). Two MLGs of temporary designated groups AF1 and PK1, first detected in Afghanistan, 2012 (MLG #103) and Pakistan, 2017 (MLG #185) were observed in East Africa (Kenya 2017 and Ethiopia 2022), respectively (XI, XII, Table 1). Dispersal event XI was confirmed by a shared virulence phenotype in source and recipient areas.

## Discussion

The present study represents one of the largest and most comprehensive spatiotemporal surveys of a major crop pathogen to date (cf. Savary et al. (2019)). Sampling covered major wheat growing regions in 41 countries on six continents, i.e., regular sampling in Africa, Asia, Europe, and South America, was connected with long-term surveillance activities in Australia through key reference samples from first yellow rust appearance on the continent and until recent time (1978-2021). North American samples were fewer and more sporadic, e.g., from the historic Stubbs collection (Thach et al., 2015), epidemiological studies on pathogen adaptation to warmer temperatures and host resistance (Milus et al., 2006; Milus et al., 2009; Milus et al., 2015b), as well as two reference samples from CIMMYT (Mexico) and two from a recent Canadian study (Holden et al., 2025). Altogether, this allowed us to explore fundamental hypotheses about evolutionary origin and LDD dispersal capacity of novel and highly significant clonal groups within *Pst*, considering current insights about pathogen diversity and dynamics at regional and global scales. This also includes comparative analyses of limited diversity in clonal populations in Europe, the Americas, Australia, Africa, the Middle East and West Asia (Hovmøller et al., 2002; Markell and Milus, 2006; Hovmøller et al., 2008; Hubbard et al., 2015; Thach et al., 2016; Ali et al., 2017b; Ding et al., 2021) versus high diversity in the near Himalayan region including China, Pakistan, Nepal and Bhutan (Duan et al., 2010; Ali et al., 2014a; Hovmøller et al., 2016; Thach et al., 2016; Khan et al., 2019). These studies in clonal as well as recombinant populations took advantage of almost identical genotyping methodology and procedures (Ali et al., 2017a; Thach et al., 2025), and revealed very different genotype profiles and multiple cases of unique allele sizes for high-diversity populations in the near Himalayan region and clonal populations elsewhere.

### Somatic hybridization by nuclear swap between parental pairs of isolates

Our study presents two sets of independent data which document for the first time in yellow rust populations the natural emergence of somatic hybrids by nuclear reassortment between co-existing pathogen individuals. Nuclear haplotype A of a prevalent European race (a.k.a. Solstice-Oakley) in PstS0 (MLG #280) provided an almost 100% match with a nucleus haplotype of the founder MLG (#244) of PstS10, whereas the second haplotype (B) provided a similar match with a nuclear haplotype represented by a race (a.k.a. Warrior) of the most prevalent MLG in PstS7 (#321) in the study. The SSR results were consistent with this conclusion by providing a 100% match between corresponding SSR haplotypes of the considered parental isolates and the hybrid. The only possible alternative hypothesis for emergence of a novel clonal group (in this case PstS10) with its unique combination of SSR allele sizes, would be hybridization by sexual reproduction involving the same parents. However, this scenario would be highly unlikely since *Pst* has only rarely been detected on an alternate (sexual) host, and not outside China (Berlin et al., 2013; Wang et al., 2016; Rodriguez-Algaba et al., 2022); further, PstS0 isolates are unlikely to produce fertile telia (Ali et al., 2010); and finally the fact that random mating via sexual reproduction would not result in a progeny isolate (in this case PstS10) consisting of two haplotypes identical to those observed in parental isolates due to meiotic recombination during the telial stage.

Interestingly, our SSR data also suggests that PstS13 and PstS14 emerged by nuclear reassortment between co-existing parental pairs of isolates, initially suggested by compatible allele sizes (Table S3). The existence of exclusive and shared SSR alleles for isolates of specific pairs of clonal groups further support this hypothesis. With respect to PstS13, these isolates uniquely shared allele size 172 (RJN11) with PstS4 isolates (except for a single PstS1_2 isolate from Kenya (2014)), and allele size 272 (RJO24) was exclusive for the same two groups when comparing isolates of all named clonal groups, except for three PstS3 isolates and a unique triticale isolate from Denmark (2016). In case of isolates of PstS14, allele size 179 (RJO21) was shared with PstS1_2 isolates, except for a single isolate from Syria (2009). Based on allele size compatibility and epidemiological observations, we therefore hypothesized that PstS13 is a somatic hybrid emerging from a nuclear reassortment between parental isolates of PstS7 and PstS4, whereby PstS7 was frequently detected on both wheat and triticale in northern Europe (2011-2014) and PstS4 was mainly detected on Triticale and rarely bread wheat (2008-2014) (https://agro.au.dk/forskning/internationale-platforme/wheatrust/yellow-rust-tools-maps-and-charts/genetic-groups-frequency-map).

An overlay of the PstS7 SSR haplotypes on the PstS13 genotype showed that haplotype B was consistent with our hypothesis, whereas haplotype A was excluded due to multiple allele size mismatches (data not shown). By comparing the PstS13 genotype and haplotype B of PstS7, a hypothetical PstS4 haplotype was derived, which was subsequently validated against the most prevalent SSR genotype in PstS4 (MLG #38, 72% of samples), resulting in a 100% allelic match based on the 16 primer pairs that produced a positive genome blast result (Table S5). This suggests that the hybridization resulting in PstS13 (MLG #21) most likely took place on Triticale, however, resulting in a significant impact on *Pst* epidemiology on a wider panel of crops, including durum wheat and spring wheat in particular (Anibal Carmona et al., 2019; Ding et al., 2021).

Similarly, we hypothesized that PstS14 is a somatic hybrid emerging from a nuclear reassortment between parental isolates of PstS7 and PstS1_2, which co-existed in Northwest Africa prior to 2015, where PstS14 was first detected . Interestingly, PstS14 made up 100% of samples in surveys in three consecutive years in NW Africa, i.e., 2016 (six samples), 2017 (38 samples) and 2018 (54 samples), but only observed sporadically elsewhere. Also in this case, the allele sizes of PstS7-haplotype B was a plausible component of the hybridization, and the hypothetical PstS1_2 haplotype was derived from the PstS14 genotype. Validation against the most common MLG in PstS1_2 (MLG #294, 47.9% of samples) resulted in a 100% allelic match (Table S5), and confirmed by blasting of SSR primer sequences to the chromosome scale assembly of a PstS1 representative isolate (Schwessinger et al., 2022). A schematic representation of the hybridization events resulting in PstS10, PstS13 and PstS14 are summarized in Fig. 6.

**Fig. 6.**
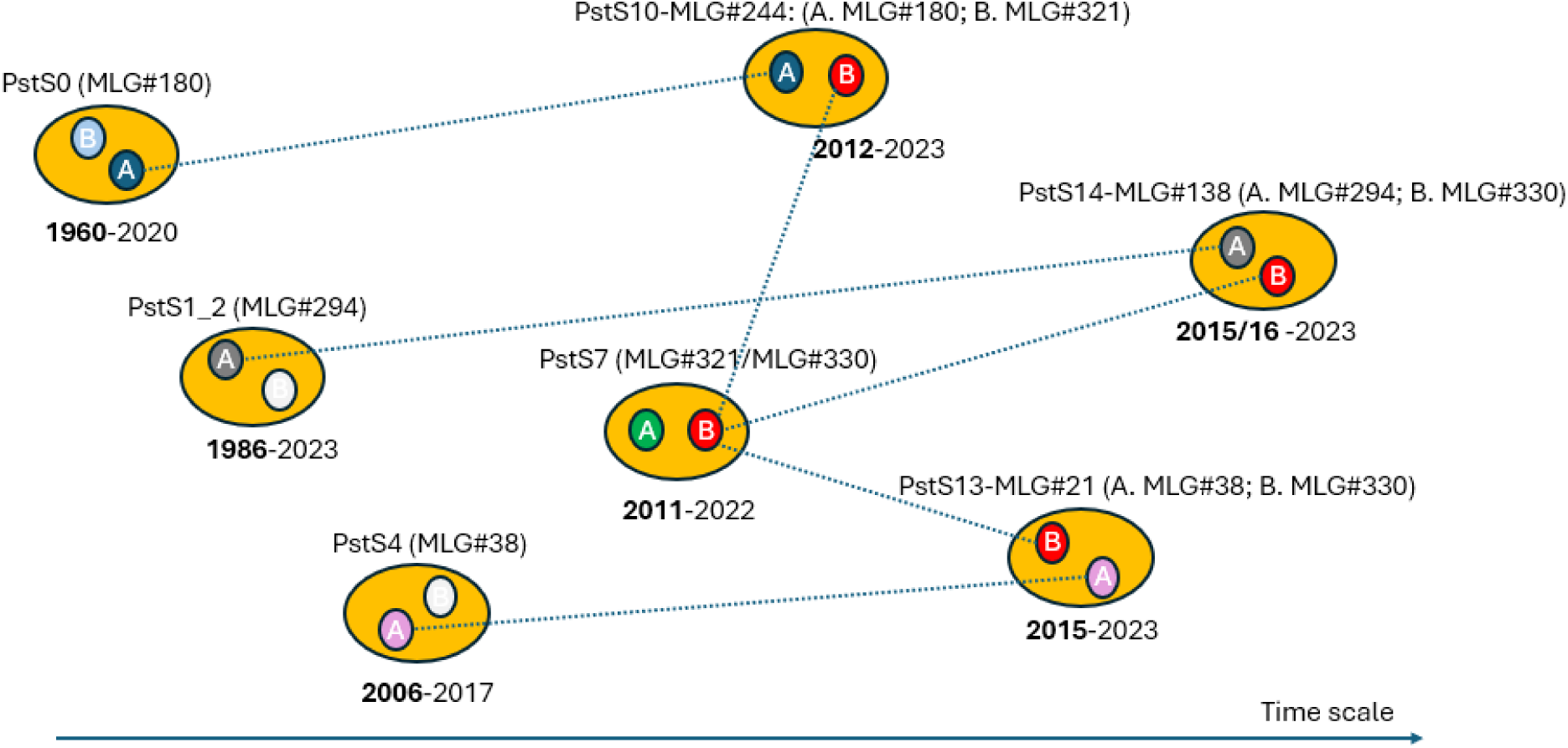
Schematic representation of three independent hybridization events supported by observed multilocus genotypes and derived nuclear haplotypes (Table S4). PstS10 was first detected in northern Europe (2012) in an area where the respective MLGs of PstS0 (#280) and PstS7 (#321) were prevalent on shared hosts of bread wheat. PstS13 was first detected in 2015, where a single MLG (#21) was detected at multiple locations in Europe, often on Triticale and rarely on bread wheat. The involved MLG of PstS4 (#38) was mainly detected on Triticale, while the MLG of PstS7 (#330) was detected widespread on triticale as well as bread wheat. PstS14 was first detected in Europe in 2015/ NW Africa 2016 (not surveyed in 2015), the involved MLGs of PstS7 (#330) and PstS1_2 (#294) being prevalent in southern Europe/NW Africa at that time. First and last sampling year of individual genetic groups in the dataset indicated (years before 2009 represented by historic reference samples).

These conclusions are consistent with the preliminary reporting of likely asexual hybridization in *Pst* in North America, involving a PstS1 isolate and a 2^nd^ unknown parent that may have resulted in PstS18 (Holden et al., 2025). PstS18 was defined recently by Thach et al. (2025) with the earliest PstS18 samples originating from five different locations in Arkansas (USA) in 2013 (Milus et al., 2015b), followed by two samples provided by CIMMYT in 2023. Interestingly, our SSR haplotype data confirm PstS1 (denoted PstS1_2 in current study) as donor of one of the nuclei in PstS18 (Table S5). This is further supported by the presence of a unique diagnostic SCAR marker for PstS1 (Walter et al., 2016). The SSR data from a different US sample revealed a complete match of unique allele sizes that can identify the donor of the 2^nd^ parental nucleus of PstS18. This isolate was first detected in 2007 according to Milus et al. (2015b). All together, these observations suggest that PstS18 indeed is a somatic hybrid emerging from two parental isolates that were already present in the USA in 2013, i.e., at the time of first documented case of PstS18.

Our results represent previously undocumented aspects of yellow rust epidemiology suggesting that somatic hybridization may be a more frequent source of agriculturally-relevant genetic diversity in wheat yellow rust than sexual recombination. Unlike other cereal rust fungi, somatic hybridization in *Pst* involving nuclear reassortment has only been reported under experimental conditions. Already in the 1960s, mixtures of urediniospore isolates of different races suggested that somatic hybridization by fusion of hyphae and subsequent nuclear reassortment was the likely source of new race variants (Little and Manners, 1967; Little and Manners, 1969). More recently, cross-infection studies resulting in 68 experimental isolates supported the occurrence of nuclear reassortment in one case, whereas most results were explained by assuming additional chromosome reassortment and crossovers (Lei et al., 2017). Earlier, Park and Wellings (2012) concluded that somatic hybridization was likely in many rust species and *formae speciales*, the process being observed in nature within and between rust species on *Linum*, *Populus*, *Senecio*, several grass species, and leaf rust on wheat. However, the exact mechanisms are generally not well understood.

### Long distance dispersal events

The potential impact of rare events of long-distance dispersal (LDD) of plant pathogens on plant health has been known for decades, e.g., Brown and Hovmøller (2002), Smart and Fry (2001), Ristaino et al. (2021), Jones (2021). The focus has often been on high impact examples, e.g. overview for *Pst* on wheat by Jin et al. (2020), rather than cases with insignificant impact, which may be overlooked and thereby contribute to an underestimation of the true frequency of LDD in crop pathogens. In this study, hypothesized migration events at the intercontinental scale (LDD) were investigated by analyses of first detection of new clonal groups and the distinct MLGs within them in eight geographically separated populations.

In seven cases, these LDD events resulted in severe and widespread disease epidemics in recipient areas, e.g., PstS13 in South America (Anibal Carmona et al. (2019), Riella et al. (2024); and PstS10 and PstS13 in Australia (https://groundcover.grdc.com.au/weeds-pests-diseases/diseases/stripe-rust-incursions-create-huge-challenges; Ding et al. (2021)), whereas five cases had only minor epidemiological impact (i.e., rare detection of corresponding clonal groups in following years). Our conclusions concerning the hypotheses of LDD versus an alternative hypothesis of hybridization of “local” isolates were supported by lack of detection of matching parental isolates in recipient areas, e.g., PstS7 in Australia and PstS1_2 and PstS4 in South America, despite extensive and widespread surveillance efforts (www.wheatrust.org; (Anibal Carmona et al., 2019; Ding et al., 2021; Riella et al., 2024)).

In four high-impact and three low-impact cases, the conclusion of very recent dispersal events was supported by identical virulence phenotypes in source and recipient areas based on assays comprising wheat experimental lines representing more than 25 different host resistance specificities (Thach et al., 2025). Because mutation from avirulence to virulence in plant pathogenic fungi, including *Pst,* may be higher than average mutation rates (Hovmøller and Justesen, 2007), and the impact likely accelerated by host induced selection (Wellings and McIntosh, 1990; Bayles et al., 2000; de Vallavieille-Pope et al., 2012a), the existence of an exact match in virulence phenotype is indicating connectivity between such populations in a short term perspective.

Considering common source and recipient areas of the 12 reported LDD events, we hypothesized seven putative transmission routes. To investigate if fungal spores could have been transported along these routes by winds, we assessed the plausibility of wind dispersal based on a combination of atmospheric trajectory simulations, high-resolution global meteorological data, data on wheat growing seasons, e.g., Bradshaw et al. (2022), literature on related fungal pathogens , and testing different spore survival times in the atmosphere (Table 2). Results indicate that both human-mediated and windborne dispersal have played an important role in intercontinental dispersal of wheat yellow rust.

**Table 2.**
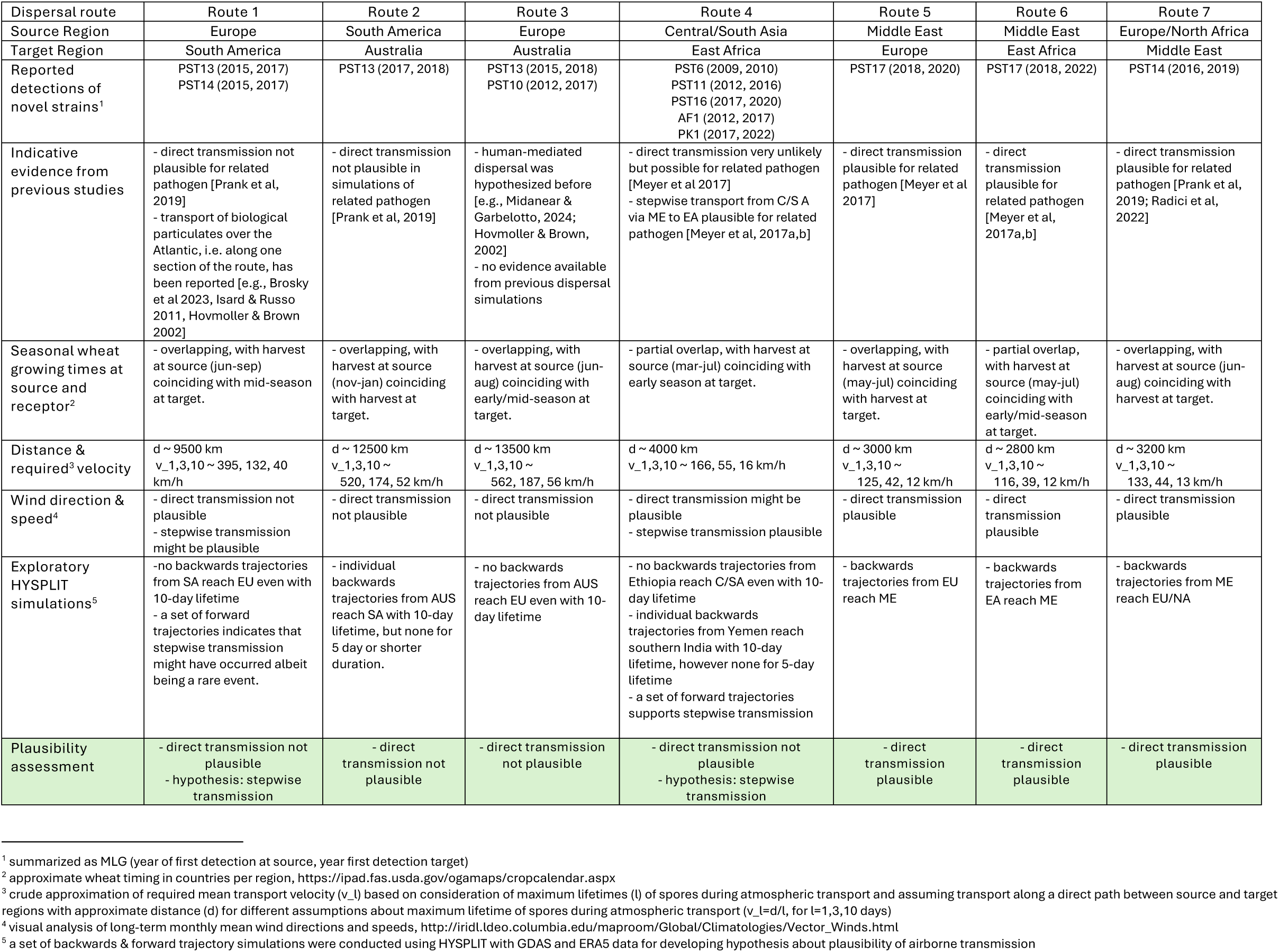
Plausibility of airborne dispersal as the causal mechanism for exotic incursions of novel Pst strains

| Dispersal route | Route 1 | Route 2 | Route 3 | Route 4 | Route 5 | Route 6 | Route 7 |
| --- | --- | --- | --- | --- | --- | --- | --- |
| Source Region | Europe | South America | Europe | Central/South Asia | Middle East | Middle East | Europe/North Africa |
| Target Region | South America | Australia | Australia | East Africa | Europe | East Africa | Middle East |
| Reported detections of novel strains<sup>1</sup> | PST13 (2015, 2017)<br>PST14 (2015, 2017) | PST13 (2017, 2018) | PST13 (2015, 2018)<br>PST10 (2012, 2017) | PST6 (2009, 2010)<br>PST11 (2012, 2016)<br>PST16 (2017, 2020)<br>AF1 (2012, 2017)<br>PK1 (2017, 2022) | PST17 (2018, 2020) | PST17 (2018, 2022) | PST14 (2016, 2019) |
| Indicative evidence from previous studies | - direct transmission not plausible for related pathogen [Prank et al., 2019]<br>- transport of biological particulates over the Atlantic, i.e. along one section of the route, has been reported [e.g., Brosky et al 2023; Isard & Russo 2011; Hovmøller & Brown 2002] | - direct transmission not plausible in simulations of related pathogen [Prank et al., 2019] | - human-mediated dispersal was hypothesized before [e.g., Midanear & Garbelotto, 2024; Hovmøller & Brown, 2002]<br>- no evidence available from previous dispersal simulations | - direct transmission very unlikely but possible for related pathogen [Meyer et al 2017]<br>- stepwise transport from C/S A via ME to EA plausible for related pathogen [Meyer et al. 2017a,b] | - direct transmission plausible for related pathogen [Meyer et al 2017] | - direct transmission plausible for related pathogen [Meyer et al., 2017a,b] | - direct transmission plausible for related pathogen [Prank et al., 2019; Radici et al., 2022] |
| Seasonal wheat growing times at source and receptor<sup>2</sup> | - overlapping, with harvest at source (jun-sep) coinciding with mid-season at target. | - overlapping, with harvest at source (nov-jan) coinciding with harvest at target. | - overlapping, with harvest at source (jun-aug) coinciding with early/mid-season at target. | - partial overlap, with harvest at source (mar-jul) coinciding with early season at target. | - overlapping, with harvest at source (may-jul) coinciding with harvest at target. | - partial overlap, with harvest at source (may-jul) coinciding with early/mid-season at target. | - overlapping, with harvest at source (jun-aug) coinciding with harvest at target. |
| Distance & required<sup>3</sup> velocity | d ~ 9500 km<br>v<sub>1,3,10</sub> ~ 395, 132, 40 km/h | d ~ 12500 km<br>v<sub>1,3,10</sub> ~ 520, 174, 52 km/h | d ~ 13500 km<br>v<sub>1,3,10</sub> ~ 562, 187, 56 km/h | d ~ 4000 km<br>v<sub>1,3,10</sub> ~ 166, 55, 16 km/h | d ~ 3000 km<br>v<sub>1,3,10</sub> ~ 125, 42, 12 km/h | d ~ 2800 km<br>v<sub>1,3,10</sub> ~ 116, 39, 12 km/h | d ~ 3200 km<br>v<sub>1,3,10</sub> ~ 133, 44, 13 km/h |
| Wind direction & speed<sup>4</sup> | - direct transmission not plausible<br>- stepwise transmission might be plausible | - direct transmission not plausible | - direct transmission not plausible | - direct transmission might be plausible<br>- stepwise transmission plausible | - direct transmission plausible | - direct transmission plausible | - direct transmission plausible |
| Exploratory HYSPLIT simulations<sup>5</sup> | - no backwards trajectories from SA reach EU even with 10-day lifetime<br>- a set of forward trajectories indicates that stepwise transmission might have occurred albeit being a rare event. | - individual backwards trajectories from AUS reach SA with 10-day lifetime, but none for 5 day or shorter duration. | - no backwards trajectories from AUS reach EU even with 10-day lifetime | - no backwards trajectories from Ethiopia reach C/SA even with 10-day lifetime<br>- individual backwards trajectories from Yemen reach southern India with 10-day lifetime, however none for 5-day lifetime<br>- a set of forward trajectories supports stepwise transmission | - backwards trajectories from EU reach ME | - backwards trajectories from EA reach ME | - backwards trajectories from ME reach EU/NA |
| Plausibility assessment | - direct transmission not plausible<br>- hypothesis: stepwise transmission | - direct transmission not plausible | - direct transmission not plausible | - direct transmission not plausible<br>- hypothesis: stepwise transmission | - direct transmission plausible | - direct transmission plausible | - direct transmission plausible |

<sup>1</sup> summarized as MLG (year of first detection at source, year first detection target)
<sup>2</sup> approximate wheat timing in countries per region, https://ipad.fas.usda.gov/ogamaps/cropcalendar.aspx <sup>3</sup> crude approximation of required mean transport velocity (v<sub>1</sub>) based on consideration of maximum lifetimes (l) of spores during atmospheric transport and assuming transport along a direct path between source and target regions with approximate distance (d) for different assumptions about maximum lifetime of spores during atmospheric transport (v<sub>1</sub>=d/l, for l=1,3,10 days)
<sup>4</sup> visual analysis of long-term monthly mean wind directions and speeds, http://iridl.ldeo.columbia.edu/maproom/Global/Climatologies/Vector_Winds.html <sup>5</sup> a set of backwards & forward trajectory simulations were conducted using HYSPLIT with GDAS and ERA5 data for developing hypothesis about plausibility of airborne transmission

Airborne dispersal appears especially plausible between the Middle East and East Africa as well as Europe. This aligns with observed, genetic similarities between *Pst* populations in the Middle East and East Africa that also indicates frequent gene flow between these regions and help explain several of the reported incursion events. The recent detections of Pst6, Pst11, Pst16, Pst17, AF1 and PK1 in East Africa after previous detections in the Middle East and Central/South Asia, provide supportive evidence for the previously proposed Rift Valley Incursion pathway for rust fungi (Meyer et al., 2017a). Direct dispersal from Central/South Asia to East Africa seems highly unlikely, but the possibility of stepwise airborne dispersal via the Middle East cannot be ruled out. Stepwise transmission is less supported by current data because PstS6 and PstS16 were not detected in the Middle East, and PstS11 appeared in the region only three years after first detection in East Africa. However, due to a relatively low sampling intensity in critical areas on the Arabian Peninsula and the Horn of Africa and simulations indicating a plausible stepwise transmission, we cannot exclude this possibility.

Direct airborne dispersal from Europe to South America or from Central/South Asia to East Africa is highly unlikely, but we hypothesize that indirect stepwise windborne transmission might have occurred, noting that a more comprehensive modelling study is required to test this. For the two dispersal events from Europe to Australia and South America to Australia, wind dispersal was not plausible, strongly indicating the role of human-mediated transport and emphasizing the importance of quantifying risks due to unintended human-mediated transmission via trade and travel.

The trajectory simulations presented here were intended for developing initial hypotheses about which dispersal mechanisms might be plausible for each incursion event. Testing as part of future work would require large-scale modelling efforts incorporating detailed spore dispersion simulations that include environmental and epidemiological conditions at source and target regions, along with sensitivity analysis covering spore survival times, release times during sporulation and locations. To advance understanding and capacity to predict global scale dispersal risks, various components are required, most notably (i) the more detailed experimental data on spore survival times during atmospheric transport obtained from large samples of viable spores, to estimate more reliably the fraction of spores remaining viable, out of extremely large numbers released during outbreaks, and (ii) a continued extension of global surveillance networks.

Altogether, our study has documented a high degree of connectivity world-wide between populations of one the most epidemic crop pathogens, yellow rust infecting cereals and grasses. Incursions from the Himalayas and elsewhere into Europe and the Mediterranean basin 10-20 years ago (Hubbard et al., 2015; Hovmøller et al., 2016; Walter et al., 2016) hybridized with isolates of pre-existing genetic groups, e.g., PstS0, PstS1_2, PstS4, the new hybrids to a large extent replaced existing population(s), spreading onwards to new continents (Hovmøller et al., 2023), where they rapidly became dominant (Anibal Carmona et al., 2019; Ding et al., 2021; Riella et al., 2024). The connectivity between populations of airborne pathogens is probably particularly evident in global cropping systems like wheat, which is grown on six continents.

The rapid raise of hybrids such as PstS10, PstS13 and PstS14, which to a large extent replaced pre-existing pathogen populations in Europe, Australia and South America, suggests that new epidemiological insights into the actual mechanisms of hybridization at both cellular and genetic levels are required. While not all hybridization events necessarily lead to increased fitness or persistence, the widespread success of these particular hybrids suggests that these may be potentially superior in fitness to their parents taking the entire epidemiological disease cycle into account, which may involve changed interactions with various components and mechanisms of host resistance.

Our data suggest that the risks and pace of pathogen evolution driven by mutation and somatic hybridization may correlate positively with the size and areas affected by ongoing epidemics. In turn, this may influence the probabilities of further hybridization and risks of unintentional pathogen spread at local, regional and global levels. Thus, there is an imminent and urgent need to sustain pathogen surveillance to better anticipate and prevent future epidemics and to facilitate development of resistant wheat cultivars. In parallel, greater awareness is needed around the risks of unintentional crop pathogen spread via human activity to enhance food security.

## Supporting information

Supplementary information part I

Supplementary information part II

## Acknowledgements

This research was supported by multiple funding sources, including a series of grants by the Bill and Melinda Gates Foundation, the UK Foreign, Commonwealth & Development Office (INV-048345, DEWAS; INV-003439, AGG; ID 0PP1133199, DGGW; OPPGD1389: DRRW), the Strategic Danish Research Council (grant no. 11-116241, RUSTFIGHT), the Danish Ministry of Food, Agriculture and Fisheries (grant no. 34120902567), Novo Nordisk Foundation (grant no. 0056457); a series of grants from Jordbruksverket (Sweden), agreement no. 21-11857/09, 25-6922/10, 25-2363/12, 4.4.11-1615/14, 4.4.11-01001/18, 4.4.11-17402/2021, Research Council of Norway (Hveterust, No. 301835), as well as long-term support by Aarhus University. Mareike Möller was supported by EU Marie Skłodowska-Curie fellowship (R-evolution, grant no. 10119509). The work of Marcel Meyer was funded by the Deutsche Forschungsgemeinschaft (DFG, German Research Foundation) – grant no. 529743941. Work at ANU was supported by DP230100941 from the Australian Research Council. Rita Tam was supported by a Grains Research and Development Corporation Graduate Research Scholarship. Rust sampling undertaken by Turkey-ICARDA Regional Cereal Rust Research Center was additionally supported by TAGEM (General Directorate of Agricultural Research and Policies of Turkey), CRP Wheat (CIMMYT), and FAO-Turkey. The authors are grateful to Ellen Jørgensen, Janne Holm Hansen, Jakob Sørensen, and Steen Meier, Aarhus University, for technical assistance during yellow rust genotyping and virulence phenotyping of 1000s of rust isolates at the Global Rust Reference Center during 15 years, Sambasivam Periyannan, Andrew Milgate, Ashley Jones, Zhenyan Luo, Zhara Chew, Eric Pereira and Danish Baig for sample characterization and assistance in the genomic research concerning isolates of PstS0, PstS7 and PstS10 (ANU), and Ezgi Kurtulus, Handan Kavas and Ali Kadiroğlu for assisting in collection and preparing samples provided by the Turkey-ICARDA Regional Cereal Rust Research Center. The deposition and access to a reference collection of *Pst* isolates from USA up to 2015 by Eugine Milus is highly appreciated, as well as the contributions of ∼ 420 people assisting in disease surveillance, sampling and submission of rust samples throughout the period 2009-2023, often under difficult and challenging conditions, without whom this work would not have been possible (for details, see Table S6).

## Author contributions

DH and MSH conceived and designed the surveillance program; TT carried out the experimental SSR genotyping work under the supervision of AFJ, including data alignment and quality check and population genetic analyses; VR assisted in genotyping of samples from South America; JRA performed virulence phenotyping of putative migrant isolates and assisted in SSR genotyping and data analyses; JGH developed the web-based data management and display system for world-wide prevalence of *Pst* genetic groups; KN supplied raw SSR data for Middle East samples 2018-2020; RP was curator of reference samples representing the Australian *Pst* population 1979-2021, undertook race analysis, and was responsible for associated funding; RT and MM performed the sequencing and genomic data analyses of PstS10 hybrid and parental isolates; BS and JR contributed to genomic data analysis, supervised and secured funding for the genomic work and provided extensive feedback on the manuscript; PS RT and MM performed the sequencing and genomic data analyses of PstS10 hybrid and parental isolates; PS supervised the PhD work of VR in South America; MME conducted trajectory simulations of the plausibility of wind dispersal of spores based on meteorological datasets; MSH was curator of samples hosted by the GRRC, supervised project activities anchored at GRRC, wrote drafts of the manuscript; all authors revised and accepted the final manuscript.

## Data availability

All raw sequencing data generated in this study have been submitted to NCBI BioProject database under accession number PRJNA1256629. The SSR genotypic data of the 3240isolates included in the present study are available at https://doi.org/10.5281/zenodo.15481965

