## Supplementary information part I for "Long-term surveillance reveals hybridization by nuclear reassortment and intercontinental spread as major evolutionary drivers in wheat yellow rust"

Article acceptance date: tbd

The following Supporting Information is available for this article:

**Fig. S1** Geographical sampling areas and number of samples per country across 41 countries and six continents.

**Fig. S2** Genotypic resolution based on 18 polymorphic SSR markers with assigned chromosomal location.

**Fig. S3** Approach and hierarchy of genetic information for exploring hypotheses of hybridization events in *Pst*. Initial focus on three novel, clonal groups that were first detected in the dataset in 2012 (PstS10), 2015 (PstS13) and 2015/16 (PstS14), respectively.

**Table S1** Worldwide collection of 3240 P. striiformis samples from 41 countries and six continents grouped according to geographical sampling origin. 3180 samples were collected in the core study period (2009-2023) whereas 60 samples were included as reference samples mainly from Australia and North America, subsets of these collected before 2009.

**Table S2** SSR diversity in P. striiformis clonal groups defined according to Thach et al. (2025). Unique MLG identification numbers with reference to hybridiaztion and LDD events shown in right column.

**Table S3** Number of allele size mismatches at 19 SSR loci for parental isolates of clonal groups hypothesized to result in PstS10, PstS13 and PstS14, respectively (year of first detection in parenthesis). Parental variant MLGs detected prior to first detection of hypothesized hybrid, and resampled at least once, were considered in the initial screen. Intervals reflect presence of multiple MLGs in considered parental group.

**Table S4** Calculated amplicon length and chromosome location of SSR primers based on haplo-phased genome information of representative PstS0, PstS7 and PstS10 isolates. The allele sized considered in the study and scored for individual samples are indicated.

**Table S5** Calculated and scored allele sizes of haplotypes of putative parental and resulting hybrids represented by PstS13, PstS14 and PstS18 (n.s.: not scored)

**Table S6** Overview of 419 people contributing to world-wide sampling of yellow rust between 2009 and 2023. When available, affiliations at time of sampling have been provided.

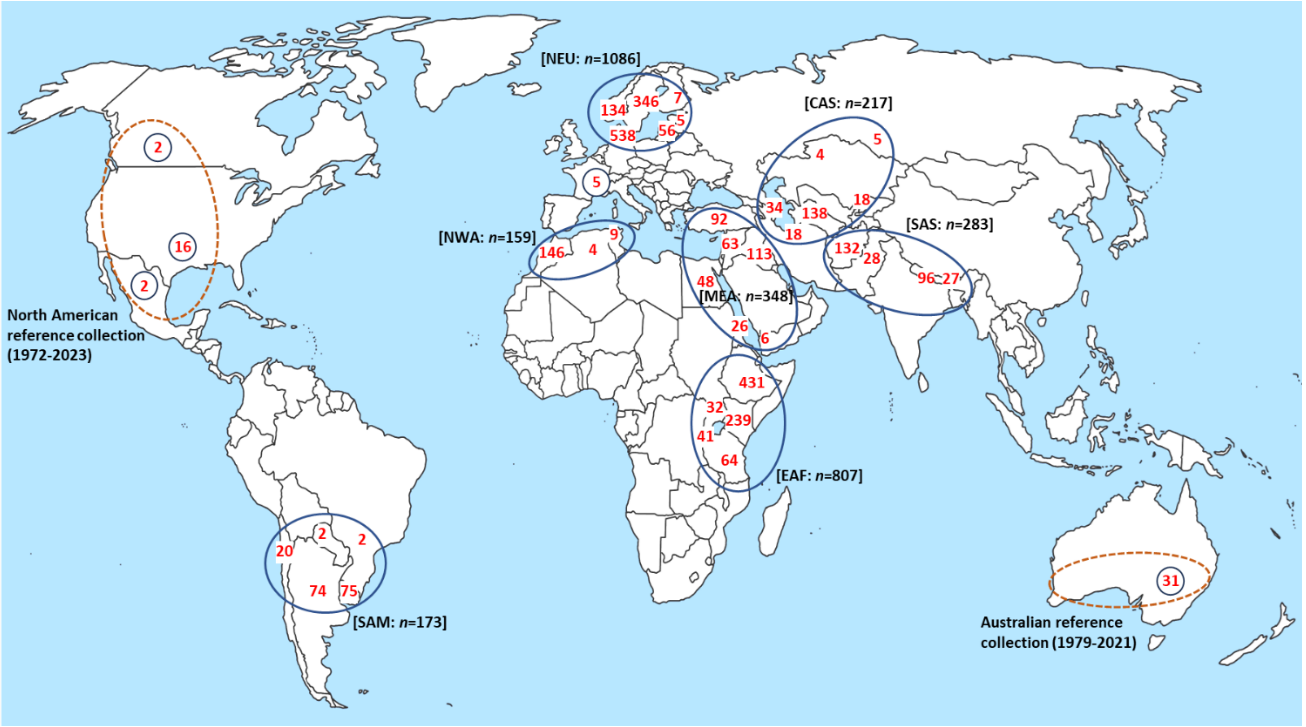


**Fig. S1** Geographical sampling areas and number of samples per country across 41 countries and six continents.

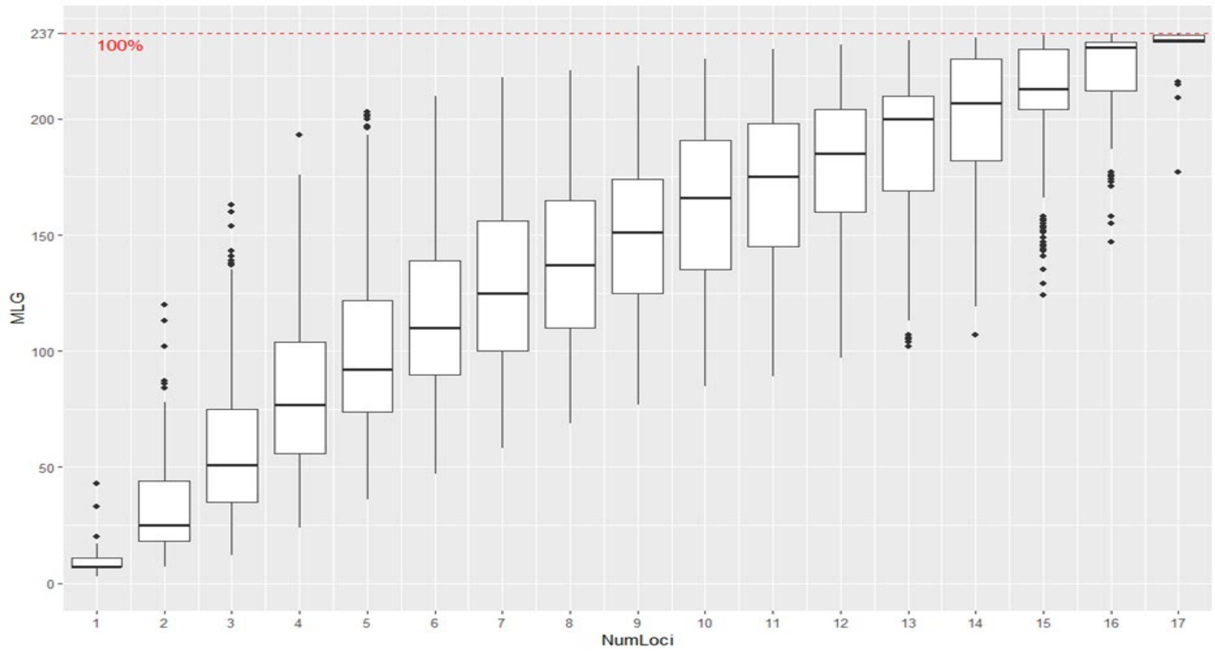


**Fig. S2** Genotypic resolution using 18 polymorphic SSR markers, where the X axis represents number of loci (minus 1) and the Y axis number of observed MLGs (*n*= 237). The bars reflect median, upper and lower quartiles, whiskers and outliers.

***1. Validation of allele sizes of MLGs representing novel groups with those of co-existing clonal groups in the dataset***

⇨ MLGs with allele sizes (including null alleles) that were not represented in the founder MLG of a novel group were excluded as potential parents in hypothesized hybridization by nuclear reassortment (Table S3).

⬇

**2. Determination of nuclear-specific SSR haplotypes and genomic validation**

a. SSR haplotypes of hypothesized parental isolates (PstS0 and PstS7) contributing to PstS10 (according to step 1) determined by blast analyses of SSR primer sequence pairs on genomic data, resulting in SSR location on haplo-phased genome maps of PstS0, PstS7 and PstS10, followed by inference of SSR haplotypes of observed MLGs and variants hereof.
b. Validation of nuclear reassortment by SNP counts and average identity based on pairwise genome alignments (Fig. 4).

⬇

**3. SSR haplotype inference of parental pair resulting in the PstS14 hybrid**

a. Following step 1-2, PstS1_2 and PstS7 were tested as potential parents of founder MLG of PstS14 (cf. Table S3). Haplotype A of PstS7 excluded due to mismatching allele sizes. The genotype of PstS14 and haplotype B of PstS7 revealed the 2nd haplotype of PstS14 (likely originating from PstS1_2).
b. Validation: Haplo-phased genome information and derived SSR allele sizes confirmed PstS1_2 as one of the parents (data not shown). Both parental haplotypes that resulted in 100% match of the observed founder MLG of PstS14 were confirmed to co-existed in NW Africa 2013-2014 co-existed in NW Africa 2013-2014.

⬇

**4. SSR haplotype inference of parental pair resulting in the PstS13 hybrid**

a. Following step 1-2, PstS4 and PstS7 were tested as potential parents of founder MLG of PstS13 (cf. Table S3). Haplotype A of PstS7 excluded due to mismatching allele sizes. The genotype of PstS13 and haplotype B of PstS7revealed the 2nd haplotype of PstS13 (likely originating from PstS4).
b. Validation: Allele sizes of hypothesized parental haplotypes, which co-existed on Triticale in northern Europe 2011-2013, resulted in 100% match of the observed founder MLG of PstS13.

**Figure S3 .** Approach and hierarchy of genetic information for exploring hypotheses of hybridization events in *Pst*. Initial focus on three novel, clonal groups that were first detected in the dataset in 2012 (PstS10), 2015 (PstS13) and 2015/16 (PstS14), respectively.
