## Supplementary information part II for "Long-term surveillance reveals hybridization by nuclear reassortment and intercontinental spread as major evolutionary drivers in wheat yellow rust"

TABLE S1 . Worldwide collection of 3240 *P. striiformis* samples from 41 countries and six continents grouped according to geographical sampling origin. 3180 samples were collected in the core study period (2009-2023) whereas 60 samples were included as reference samples mainly from Australia and North America, subsets of these collected before 2009.

| | | | | | References | | |
|---|---|---|---|---|---|---|---|
| **Geographic sampling origin** | **Country** | **Period I (2009-2016)** | **Period II (2017-2023)** | **Grand Total (2009-2023)** | **<2009** | **2009-2023** | **Sampling years** |
| **Australasia** (AU) | Australia | 1 | 7 | 8 | 23 | 8 | 1979, 1982-1988, 1991, 1992, 1998, 1999, 2002, 2004- |
| **Australasia Total** (reference samples) | | **1** | **7** | **8** | **23** | **8** | 2009, 2017, 2018, 2020, 2021 |
| **Central_west Asia** | Azerbaijan | 31 | 3 | 34 | | | 2009, 2010, 2013, 2015-2018 |
| **(CWA)** | Iran | 8 | 10 | 18 | | | 2012, 2014, 2015, 2018, 2019 |
| | Kazakhstan | 4 | 0 | 4 | | | 2015 |
| | Russia | 0 | 5 | 5 | | | 2017 |
| | Tajikistan | 18 | 0 | 18 | | | 2010, 2011, 2013 |
| | Uzbekistan | 53 | 85 | 138 | | | 2011, 2013, 2014, 2016-2019 |
| **Central_west Asia Total** | | **114** | **103** | **217** | | | |
| **East Africa** | Ethiopia | 58 | 422 | 480 | 1 | | 2010, 2013-2023 |
| **(EAF)** | Kenya | 27 | 239 | 266 | | | 2009, 2011, 2013, 2014, 2017-2023 |
| | Rwanda | 28 | 13 | 41 | | | 2014, 2015, 2017, 2018 |
| | Tanzania | 31 | 48 | 79 | | | 2013, 2015-2018, 2021, 2022 |
| | Uganda | 0 | 48 | 48 | | | 2019, 2023 |
| **East Africa Total** | | **144** | **770** | **914** | **1** | | |
| **Europe** | France | 1 | 1 | 2 | | 2 | 2012; 2017 |
| (reference samples) | Germany | 0 | 0 | | 1 | | 1960 |
| | Greece | 0 | 0 | | 1 | | 1973 |
| | United Kingdom | 0 | 0 | | 1 | | 1975 |
| **Europe Total** | | **1** | **1** | **2** | **3** | **2** | **Refs** |
| **Middle East** | Egypt | 19 | 29 | 48 | | | 2013, 2018, 2019 |
| **(MEA)** | Eritrea | 12 | 14 | 26 | | | 2011-2013, 2016-2018 |
| | Iraq | 59 | 54 | 113 | | | 2010, 2011, 2013, 2014, 2016, 2017, 2021, 2022 |
| | Jordan | 0 | 12 | 12 | | | 2019 |
| | Lebanon | 3 | 29 | 32 | | | 2018, 2019 |
| | Syria | 10 | 9 | 19 | | | 2009, 2010, 2019 |
| | Turkey | 5 | 87 | 92 | | | 2011, 2017-2020 |
| | Yemen | 6 | 0 | 6 | | | 2014 |
| **Middle East Total** | | **114** | **234** | **348** | | | |
| **North America** | United States | 8 | 0 | 8 | 8 | 8 | 1972, 1980, 1981, 1983, 2000, 2003, 2014, 2015 |
| (reference samples) | Canada | | 2 | 2 | | 2 | 2022 |
| | Mexico | 0 | 2 | 2 | | 2 | 2023 |
| **North America Total** | | **8** | **4** | **12** | **8** | **12** | **Refs** |
| **Northern Europe** | Denmark | 204 | 334 | 538 | 3 | | 2002, 2003, 2006, 2009-2023 |
| **(NEU)** | Estonia | 0 | 5 | 5 | | | 2018, 2021 |
| | Finland | 7 | 0 | 7 | | | 2016 |
| | Latvia | 6 | 50 | 56 | | | 2015-2021 |
| | Norway | 35 | 99 | 134 | | | 2014-2023 |
| | Sweden | 163 | 183 | 346 | | | 2009-2023 |
| **Northern Europe Total** | | **415** | **671** | **1086** | **3** | | **Refs** |
| **Northwest Africa** | Algeria | 4 | 0 | 4 | | | 2014 |
| **(NWA)** | Morocco | 37 | 109 | 146 | | | 2009, 2013, 2016-2019 |
| | Tunisia | 0 | 9 | 9 | | | 2022, 2023 |
| **Northwest Africa Total** | | **41** | **118** | **159** | | | |
| **South America** | Argentina | 4 | 70 | 74 | | | 2015, 2017, 2018 |
| **(SAM)** | Brazil | 2 | 0 | 2 | | | 2010 |
| | Chile | 2 | 18 | 20 | | | 2010, 2018 |
| | Paraguay | 0 | 2 | 2 | | | 2021 |
| | Uruguay | 12 | 63 | 75 | | | 2010, 2021, 2022 |
| **South America Total** | | **20** | **153** | **173** | | | |
| **South Asia** | Afghanistan | 41 | 91 | 132 | | | 2009, 2010, 2012-2016, 2018, 2021 |
| **(SAS)** | Bhutan | 23 | 4 | 27 | | | 2012, 2013, 2015, 2018 |
| | Nepal | 4 | 92 | 96 | | | 2014, 2015, 2021-2023 |
| | Pakistan | 5 | 23 | 28 | | | 2014, 2017 |
| **South Asia Total** | | **73** | **210** | **283** | | | |
| **Reference samples** | | **10** | **12** | **22** | **38** | **22** | |
| **Total** | | **921** | **2259** | **3180** | **38** | **22** | **Grand total: 3240** |

Table S2. SSR diversity in *P. striiformis* clonal groups defined according to Thach et al. (2025). Unique MLG identification numbers with reference to hybridiaztion and LDD events shown in right column.

| Clonal group | No. of samples (n) | No. of MLGs/ clonal group | MLGs/ group (normalized by *n)* | Freq. most prevalent MLG/ group (in %) | Unique MLG identifier (reference #). Bold: most frequent within clonal group |
|---|---|---|---|---|---|
| PstS0 | 79 | 12 [1] | 0,152 | 54,4 | 276, **277**, 279, 280, 283, 306, 307 |
| PstS1_2 | 438 | 24 | 0,055 | 47,9 | 151-153, 163, 165, 166, 223, 226-228, 268, 273, 286-**294**, 299, 301, 308 |
| PstS3 | 3 | 1 | 0,333 | 100,0 | **42** |
| PstS4 | 51 | 3 | 0,059 | 72,0 | 36-**38** |
| PstS5 | 14 | 4 | 0,286 | 35,7 | 380-**383** |
| PstS6 | 46 | 4 | 0,087 | 91,3 | 5-**8** |
| PstS7 | 106 | 19 | 0,179 | 41,0 | 316, 318-**321**, 322-325, 329, 330, 332-335, 338, 339, 341, 377 |
| PstS8 | 128 | 15 | 0,117 | 51,6 | 317, 352-**358**, 362-367, 370 |
| PstS9 | 139 | 14 | 0,101 | 53,2 | 391, 393-**396**, 400-406, 409, 415 |
| PstS10 | 602 | 16 | 0,027 | 82,2 | 124, 222, 237, 238, 240-**244**, 247-249, 258, 259, 263, 311 |
| PstS11 | 454 | 22 | 0,048 | 48,5 | 47, 48, 51, 59-61, 63-65, 71, 72, 74, 76-78, 82, 85-87, 96, 97, 102 |
| PstS12 | 6 | 1 | 0,167 | 100,0 | **275** |
| PstS13 | 282 | 6 | 0,021 | 84,3 | 19-**21**, 22, 24, 27 |
| PstS14 | 239 | 6 | 0,025 | 72,0 | 132, 134, 136-**138**, 141 |
| PstS15 | 14 | 5 | 0,357 | 30,8 | **342**, 343, 345-347 |
| PstS16 | 278 | 13 | 0,047 | 70,1 | 159, 189, 190, 192, 194, 195, 202-**205**, 207, 219, 221 |
| PstS17 | 87 | 3 | 0,034 | 77,0 | **170**, 171, 175 |
| PstS18 | 7 | 1 | 0,143 | 100,0 | **906** |
| Other | 267 | 76 | 0,285 | | (seventy six) |
| Total PstS0-PstS18 | 2973 | 169 | 0,057 | - | |
| **Grand total** | 3240 | 245 | 0,076 | - | |

[1] Extended sampling period and area including reference samples from Europe, North America and Australia (1978-2021)

Table S3. Number[1] of allele size mismatches at 19 SSR loci for parental isolates of clonal groups hypothesized to result in PstS10, PstS13 and PstS14, respectively (year of first detection in parenthesis). Parental variant MLGs detected prior to first detection of hypothesized hybrid, and resampled at least once, were considered in the initial screen. Intervals reflect presence of multiple MLGs in considered parental group.

| | Novel hybrid group (year of first detection in parenthesis) | | |
|---|---|---|---|
| Potential parental group | PstS10 (2012) | PstS13 (2015) | PstS14 (2015/2016) |
| PstS0 | >= 0 | >=5 | 4 |
| PstS1_2 | >=2 | >=4 | >= 0 |
| PstS3 | >=3 | 2 | 2 |
| PstS4 | >=4 | >= 0 | 3 |
| PstS5 | 3 | >=4 | 4 |
| PstS6 | >=6 | 6 | 5 |
| PstS7 | >= 0 | >= 0 | >= 0 |
| PstS8 | >= 1 | >=2 | >=2 |
| PstS9 | 3 | 5 | 4 |
| PstS10 | - | >= 0 | >= 0 |
| PstS11 | - | >=5 | 4 |
| PstS12 | - | 7 | 5 |
| PstS13 | - | - | >= 0 |
| PstS14 | - | - | - |

[1]Numbers >0 imply that hypothesis of hybridization by nuclear reassortment is rejected, unless additional assumptions about parasexuality and crossovers are made.

**Table S4. Calculated amplicon length and chromosome location of SSR primers based on haplo-phased genome information of representative PstS0, PstS7 and PstS10 isolates. The allele sized considered in the study and scored for individual samples are indicated.**

**PstS0 (PBI accession no. 821559)** Separate Excel file: Chromosomal location

| primer_ID | chromosome | start | end | amplicon length | Scored allele size |
|---|---|---|---|---|---|
| RJN10F | chr10A | 3210421 | 3210403 | 227 | 224 |
| RJN10R | chr10A | 3210194 | 3210212 | | |
| RJN10F | chr10B | 3303603 | 3303585 | 224 | 221 |
| RJN10R | chr10B | 3303379 | 3303397 | | |
| RJN11F | chr10A | 640248 | 640266 | 177 | 176 |
| RJN11R | chr10A | 640425 | 640407 | | |
| RJN11F | chr10B | 612406 | 612388 | 179 | 178 |
| RJN11R | chr10B | 612227 | 612245 | | |
| RJN12F | chr13A | 3330499 | 3330480 | 193 | 196 |
| RJN12R | chr13A | 3330306 | 3330327 | | |
| RJN12F | chr13B | 3206207 | 3206188 | 193 | 196 |
| RJN12R | chr13B | 3206014 | 3206035 | | |
| RJN13F | chr1A | 2048776 | 2048794 | 150 | 147 |
| RJN13R | chr1A | 2048926 | 2048908 | | |
| RJN13F | chr1B | 2036154 | 2036172 | 150 | 147 |
| RJN13R | chr1B | 2036304 | 2036286 | | |
| RJN2F | chr8A | 1942933 | 1942916 | 185 | 187 |
| RJN2R | chr8A | 1942748 | 1942771 | | |
| RJN2F | chr8B | N/A | N/A | - | NULL |
| RJN2R | chr8B | N/A | N/A | | |
| RJN3F | chr7A | N/A | N/A | - | NULL |
| RJN3R | chr7A | N/A | N/A | | |
| RJN3F | chr7B | 3861836 | 3861854 | 341 | 340 |
| RJN3R | chr7B | 3862177 | 3862158 | | |
| RJN4F | chr4A | 952272 | 952254 | 245 | not scored |
| RJN4R | chr4A | 952027 | 952045 | | |
| RJN4F | chr4B | 943389 | 943371 | 245 | not scored |
| RJN4R | chr4B | 943144 | 943162 | | |
| RJN4F | chr6A | 4785642 | 4785660 | 258 | 255 |
| RJN4R | chr6A | 4785900 | 4785882 | | |
| RJN4F | chr6B | 4843444 | 4843462 | 258 | 255 |
| RJN4R | chr6B | 4843702 | 4843684 | | |
| RJN5F | chr2A | 3789873 | 3789891 | 231 | 230 |
| RJN5R | chr2A | 3790104 | 3790086 | | |
| RJN5F | chr2B | 3726862 | 3726880 | 227 | 226 |
| RJN5R | chr2B | 3727089 | 3727071 | | |
| RJN6F | chr5A | 2893120 | 2893102 | 320 | 318 |
| RJN6R | chr5A | 2892800 | 2892818 | | |
| RJN6F | chr5B | 2643170 | 2643152 | 320 | 318 |
| RJN6R | chr5B | 2642850 | 2642868 | | |
| RJN8F | chr2A | 328245 | 328227 | 307 | 307 |
| RJN8R | chr2A | 327938 | 327956 | | |
| RJN8F | chr2B | 363900 | 363882 | 307 | 307 |
| RJN8R | chr2B | 363593 | 363611 | | |
| RJN9F | chr10A | 1181201 | 1181183 | 334 | 334 |
| RJN9R | chr10A | 1180867 | 1180885 | | |
| RJN9F | chr10B | 1286507 | 1286489 | 332 | 332 |
| RJN9R | chr10B | 1286175 | 1286193 | | |
| RJO18F | chr8A | N/A | N/A | - | NULL |
| RJO18R | chr8A | N/A | N/A | | |
| RJO18F | chr8B | 2562428 | 2562445 | 359 | 357 |
| RJO18R | chr8B | 2562787 | 2562769 | | |
| RJO20F | chr17A | 309932 | 309949 | 290 | 284 |
| RJO20R | chr17A | 310222 | 310205 | | |
| RJO20F | chr17B | 268197 | 268214 | 290 | 284 |
| RJO20R | chr17B | 268487 | 268470 | | |
| RJO21F | chr5A | 2462279 | 2462298 | 169 | 170 |
| RJO21R | chr5A | 2462448 | 2462429 | | |
| RJO21F | chr5B | 2266183 | 2266202 | 169 | 170 |
| RJO21R | chr5B | 2266352 | 2266333 | | |
| RJO24F | chr14A | 268940 | 268919 | 268 | 275 |
| RJO24R | chr14A | 268672 | 268691 | | |
| RJO24F | chr14B | 242259 | 242238 | 286 | 293 |
| RJO24R | chr14B | 241973 | 241992 | | |
| RJO27F | chr5A | 2460379 | 2460360 | 230 | 232 |
| RJO27R | chr5A | 2460149 | 2460173 | | |
| RJO27F | chr5B | 2264280 | 2264261 | 228 | 230 |
| RJO27R | chr5B | 2264052 | 2264076 | | |
| RJO3F | chr10A | 212449 | 212433 | 204 | not scored |
| RJO3R | chr10A | 212245 | 212264 | | |
| RJO3F | chr10B | 205823 | 205807 | 206 | not scored |
| RJO3R | chr10B | 205617 | 205636 | | |
| RJO4F | chr7A | 2619008 | 2618991 | 198 | 202 |
| RJO4R | chr7A | 2618810 | 2618829 | | |
| RJO4F | chr7B | 2634876 | 2634859 | 195 | 199 |
| RJO4R | chr7B | 2634681 | 2634700 | | |
| WU12F | chr11A | 2883611 | 2883592 | 330 | not scored |
| WU12R | chr11A | 2883281 | 2883300 | | |
| WU12F | chr11A | 2911245 | 2911264 | 327 | 323 |
| WU12R | chr11A | 2911572 | 2911553 | | |
| WU12F | chr11B | 2903482 | 2903463 | 330 | not scored |
| WU12R | chr11B | 2903152 | 2903171 | | |
| WU12F | chr11B | 2929920 | 2929939 | 336 | 332 |
| WU12R | chr11B | 2930256 | 2930237 | | |
| WU6F | chr3A | 3045693 | 3045712 | 210 | 210 |
| WU6R | chr3A | 3045903 | 3045884 | | |
| WU6F | chr3B | 2896601 | 2896620 | 210 | 210 |
| WU6R | chr3B | 2896811 | 2896792 | | |

**PstS7 (DK09_11SP)**

| primer_ID | chromosome | start | end | amplicon length | Scored allele size |
|---|---|---|---|---|---|
| RJN10F | chr10A | 3289476 | 3289458 | 227 | 224 |
| RJN10R | chr10A | 3289249 | 3289267 | | |
| RJN10F | chr10B | 3306685 | 3306667 | 227 | 221 |
| RJN10R | chr10B | 3306458 | 3306476 | | |
| RJN11R | chr10A | 611084 | 611102 | 183 | 176 |
| RJN11F | chr10A | 611267 | 611249 | | |
| RJN11F | chr10B | 614596 | 614578 | 177 | 178 |
| RJN11R | chr10B | 614419 | 614437 | | |
| RJN12F | chr13A | 4288859 | 4288840 | 193 | 196 |
| RJN12R | chr13A | 4288666 | 4288687 | | |
| RJN12F | chr13B | 3910504 | 3910485 | 193 | 196 |
| RJN12R | chr13B | 3910311 | 3910332 | | |
| RJN13F | chr1A | 2024841 | 2024859 | 150 | 147 |
| RJN13R | chr1A | 2024991 | 2024973 | | |
| RJN13F | chr1B | 2020563 | 2020581 | 153 | 147 |
| RJN13R | chr1B | 2020716 | 2020698 | | |
| RJN2F | chr8A | N/A | N/A | - | NULL |
| RJN2R | chr8A | N/A | N/A | | |
| RJN2F | chr8B | N/A | N/A | - | NULL |
| RJN2R | chr8B | N/A | N/A | | |
| RJN3F | chr7A | 3886733 | 3886751 | 337 | 336 |
| RJN3R | chr7A | 3887070 | 3887051 | | |
| RJN3F | chr7B | 3999685 | 3999703 | 337 | 336 |
| RJN3R | chr7B | 4000022 | 4000003 | | |
| RJN4F | chr4A | 926682 | 926664 | 245 | not scored |
| RJN4R | chr4A | 926437 | 926455 | | |
| RJN4F | chr4B | 957436 | 957418 | 245 | not scored |
| RJN4R | chr4B | 957191 | 957209 | | |
| RJN4F | chr6A | 5059285 | 5059303 | 258 | 255 |
| RJN4R | chr6A | 5059543 | 5059525 | | |
| RJN4F | chr6B | 5152352 | 5152370 | 258 | 255 |
| RJN4R | chr6B | 5152610 | 5152592 | | |
| RJN5F | chr2A | 3765123 | 3765141 | 227 | 226 |
| RJN5R | chr2A | 3765350 | 3765332 | | |
| RJN5F | chr2B | 3725462 | 3725480 | 227 | 226 |
| RJN5R | chr2B | 3725689 | 3725671 | | |
| RJN6F | chr5A | 2721477 | 2721459 | 320 | 318 |
| RJN6R | chr5A | 2721157 | 2721175 | | |
| RJN6F | chr5B | 2690924 | 2690906 | 317 | 315 |
| RJN6R | chr5B | 2690607 | 2690625 | | |
| RJN8F | chr2A | 340275 | 340257 | 316 | 316 |
| RJN8R | chr2A | 339959 | 339977 | | |
| RJN8F | chr2B | 336661 | 336643 | 307 | 307 |
| RJN8R | chr2B | 336354 | 336372 | | |
| RJN9F | chr10A | 1271227 | 1271209 | 332 | 332 |
| RJN9R | chr10A | 1270895 | 1270913 | | |
| RJN9F | chr10B | 1279224 | 1279206 | 332 | 332 |
| RJN9R | chr10B | 1278892 | 1278910 | | |
| RJO18F | N/A | N/A | N/A | - | **331** |
| RJO18R | N/A | N/A | N/A | | |
| RJO18F | N/A | N/A | N/A | - | **331** |
| RJO18R | N/A | N/A | N/A | | |
| RJO20F | chr17A | 293430 | 293447 | 290 | 284 |
| RJO20R | chr17A | 293720 | 293703 | | |
| RJO20F | chr17B | 314405 | 314422 | 293 | 287 |
| RJO20R | chr17B | 314698 | 314681 | | |
| RJO21F | chr5A | 2343262 | 2343281 | 169 | 170 |
| RJO21R | chr5A | 2343431 | 2343412 | | |
| RJO21F | chr5B | 2296615 | 2296634 | 169 | 170 |
| RJO21R | chr5B | 2296784 | 2296765 | | |
| RJO24F | chr14A | 243328 | 243307 | 286 | 293 |
| RJO24R | chr14A | 243042 | 243061 | | |
| RJO24F | chr14B | 254270 | 254249 | 277 | 284 |
| RJO24R | chr14B | 253993 | 254012 | | |
| RJO27F | chr5A | N/A | N/A | - | NULL |
| RJO27R | chr5A | N/A | N/A | | |
| RJO27F | chr5B | N/A | N/A | - | NULL |
| RJO27R | chr5B | N/A | N/A | | |
| RJO3F | chr10A | 198079 | 198063 | 204 | not scored |
| RJO3R | chr10A | 197875 | 197894 | | |
| RJO3F | chr10B | 199074 | 199058 | 206 | not scored |
| RJO3R | chr10B | 198868 | 198887 | | |
| RJO4F | chr7A | 2667464 | 2667447 | 195 | 199 |
| RJO4R | chr7A | 2667269 | 2667288 | | |
| RJO4F | chr7B | 2684339 | 2684322 | 195 | 199 |
| RJO4R | chr7B | 2684144 | 2684163 | | |
| WU12F | chr11A | 2926322 | 2926303 | 330 | not scored |
| WU12R | chr11A | 2925992 | 2926011 | | |
| WU12F | chr11A | 2952735 | 2952754 | 336 | 332 |
| WU12R | chr11A | 2953071 | 2953052 | | |
| WU12F | chr11B | 2932847 | 2932828 | 330 | not scored |
| WU12R | chr11B | 2932517 | 2932536 | | |
| WU12F | chr11B | 2959260 | 2959279 | 336 | 332 |
| WU12R | chr11B | 2959596 | 2959577 | | |
| WU6F | chr3A | 2932792 | 2932811 | 210 | 210 |
| WU6R | chr3A | 2933002 | 2932983 | | |
| WU6F | chr3B | 2918940 | 2918959 | 210 | 210 |
| WU6R | chr3B | 2919150 | 2919131 | | |

**PstS10 (PBI accession no. 221731)**

| primer_ID | chromosome | start | end | amplicon length | Scored size |
|---|---|---|---|---|---|
| RJN10F | chr10A | 3199965 | 3199947 | 227 | 224 |
| RJN10R | chr10A | 3199738 | 3199756 | | |
| RJN10F | chr10B | 3303093 | 3303075 | 227 | 221 |
| RJN10R | chr10B | 3302866 | 3302884 | | |
| RJN11F | chr10A | 632653 | 632671 | 177 | 176 |
| RJN11R | chr10A | 632830 | 632812 | | |
| RJN11F | chr10B | 613239 | 613221 | 177 | 178 |
| RJN11R | chr10B | 613062 | 613080 | | |
| RJN12F | chr13A | 3375688 | 3375669 | 193 | 196 |
| RJN12R | chr13A | 3375495 | 3375516 | | |
| RJN12F | chr13B | 3392365 | 3392346 | 193 | 196 |
| RJN12R | chr13B | 3392172 | 3392193 | | |
| RJN13F | chr1A | 2055839 | 2055857 | 150 | 147 |
| RJN13R | chr1A | 2055989 | 2055971 | | |
| RJN13F | chr1B | 2029078 | 2029096 | 153 | 147 |
| RJN13R | chr1B | 2029231 | 2029213 | | |
| RJN2F | chr8A | 1970887 | 1970870 | 187 | 187 |
| RJN2R | chr8A | 1970700 | 1970723 | | |
| RJN2F | chr8B | N/A | N/A | - | NULL |
| RJN2R | chr8B | N/A | N/A | | |
| RJN3F | chr7A | N/A | N/A | - | NULL |
| RJN3R | chr7A | N/A | N/A | | |
| RJN3F | chr7B | 3922332 | 3922350 | 337 | 336 |
| RJN3R | chr7B | 3922669 | 3922650 | | |
| RJN4F | chr4A | 958583 | 958565 | 245 | not scored |
| RJN4R | chr4A | 958338 | 958356 | | |
| RJN4F | chr4B | 956798 | 956780 | 245 | not scored |
| RJN4R | chr4B | 956553 | 956571 | | |
| RJN4F | chr6A | 4871451 | 4871469 | 258 | 255 |
| RJN4R | chr6A | 4871709 | 4871691 | | |
| RJN4F | chr6B | 4878859 | 4878877 | 258 | 255 |
| RJN4R | chr6B | 4879117 | 4879099 | | |
| RJN5F | chr2A | 3782817 | 3782835 | 231 | 230 |
| RJN5R | chr2A | 3783048 | 3783030 | | |
| RJN5F | chr2B | 3723966 | 3723984 | 227 | 226 |
| RJN5R | chr2B | 3724193 | 3724175 | | |
| RJN6F | chr5A | 2883599 | 2883581 | 320 | 318 |
| RJN6R | chr5A | 2883279 | 2883297 | | |
| RJN6F | chr5B | 2690470 | 2690452 | 317 | 315 |
| RJN6R | chr5B | 2690153 | 2690171 | | |
| RJN8F | chr2A | 327281 | 327263 | 307 | 307 |
| RJN8R | chr2A | 326974 | 326992 | | |
| RJN8F | chr2B | 335174 | 335156 | 307 | 307 |
| RJN8R | chr2B | 334867 | 334885 | | |
| RJN9F | chr10A | 1173648 | 1173630 | 334 | 334 |
| RJN9R | chr10A | 1173314 | 1173332 | | |
| RJN9F | chr10B | 1275452 | 1275434 | 332 | 332 |
| RJN9R | chr10B | 1275120 | 1275138 | | |
| RJO18F | N/A | N/A | N/A | - | **337** |
| RJO18R | N/A | N/A | N/A | | |
| RJO18F | N/A | N/A | N/A | - | **331** |
| RJO18R | N/A | N/A | N/A | | |
| RJO20F | chr17A | 48111 | 48128 | 290 | 284 |
| RJO20R | chr17A | 48401 | 48384 | | |
| RJO20F | chr17B | 44737 | 44754 | 293 | 287 |
| RJO20R | chr17B | 45030 | 45013 | | |
| RJO21F | chr5A | 2466815 | 2466834 | 169 | 170 |
| RJO21R | chr5A | 2466984 | 2466965 | | |
| RJO21F | chr5B | 2296134 | 2296153 | 169 | 170 |
| RJO21R | chr5B | 2296303 | 2296284 | | |
| RJO24F | chr14A | 268998 | 268977 | 268 | 275 |
| RJO24R | chr14A | 268730 | 268749 | | |
| RJO24F | chr14B | 252849 | 252828 | 277 | 284 |
| RJO24R | chr14B | 252572 | 252591 | | |
| RJO27F | chr5A | 2464915 | 2464896 | 230 | 232 |
| RJO27R | chr5A | 2464685 | 2464709 | | |
| RJO27F | chr5B | N/A | N/A | - | NULL |
| RJO27R | chr5B | N/A | N/A | | |
| RJO3F | chr10A | 204851 | 204835 | 204 | not scored |
| RJO3R | chr10A | 204647 | 204666 | | |
| RJO3F | chr10B | 197682 | 197666 | 206 | not scored |
| RJO3R | chr10B | 197476 | 197495 | | |
| RJO4F | chr7A | 2603871 | 2603854 | 198 | 202 |
| RJO4R | chr7A | 2603673 | 2603692 | | |
| RJO4F | chr7B | 2683457 | 2683440 | 195 | 199 |
| RJO4R | chr7B | 2683262 | 2683281 | | |
| WU12F | chr11A | 2874759 | 2874740 | 330 | not scored |
| WU12R | chr11A | 2874429 | 2874448 | | |
| WU12F | chr11A | 2902391 | 2902410 | 327 | 323 |
| WU12R | chr11A | 2902718 | 2902699 | | |
| WU12F | chr11B | 2933066 | 2933047 | 330 | not scored |
| WU12R | chr11B | 2932736 | 2932755 | | |
| WU12F | chr11B | 2959480 | 2959499 | 336 | 332 |
| WU12R | chr11B | 2959816 | 2959797 | | |
| WU6F | chr3A | 3072874 | 3072893 | 210 | 210 |
| WU6R | chr3A | 3073084 | 3073065 | | |
| WU6F | chr3B | 2918570 | 2918589 | 210 | 210 |
| WU6R | chr3B | 2918780 | 2918761 | | |

**Table S5. Calculated and scored allele sizes of haplotypes of putative parental and resulting hybrids represented by PstS13, PstS14 and PstS18 (n.s.: not scored)**

| | primer_ID | RJN10 | | RJN11 | | RJN12 | | RJN13 | | RJN2 | | RJN3 | | RJN4 | | RJN4 | | RJN5 | | RJN6 | | RJN8 | | RJN9 | | RJO18 | | RJO20 | | RJO21 | | RJO24 | | RJO27 | | RJO3 | | RJO4 | | WU12 | | | | WU6 | |
|---|---|---|---|---|---|---|---|---|---|---|---|---|---|---|---|---|---|---|---|---|---|---|---|---|---|---|---|---|---|---|---|---|---|---|---|---|---|---|---|---|---|---|---|---|---|
| | chromosome | chr10A | chr10B | chr10A | chr10B | chr13A | chr13B | chr1A | chr1B | chr8A | chr8B | chr7A | chr7B | chr4A | chr4B | chr6A | chr6B | chr2A | chr2B | chr5A | chr5B | chr2A | chr2B | chr10A | chr10B | chr8A | chr8B | chr17A | chr17B | chr5A | chr5B | chr14A | chr14B | chr5A | chr5B | chr10A | chr10B | chr7A | chr7B | chr11A | chr11B | chr11A | chr11B | chr3A | chr3B |
| **PstS7 (DK09_11SP)** | Calculated size (genomic data) | 227 | 227 | 183 | 177 | 193 | 193 | 150 | 153 | - | - | 337 | 337 | 245 | 245 | 258 | 258 | 227 | 227 | 320 | 317 | 316 | 307 | 332 | 332 | - | - | 290 | 293 | 169 | 169 | 286 | 277 | - | - | 204 | 206 | 195 | 195 | 330 | 330 | 336 | 336 | 210 | 210 |
| | Scored size (MLG #321) | 224 | 224 | 182 | 176 | 196 | 196 | 147 | 150 | NULL | NULL | 336 | 336 | n.s. | n.s. | 255 | 255 | 226 | 226 | 318 | 315 | 316 | 307 | 332 | 332 | **331** | **331** | 284 | 287 | 170 | 170 | 293 | 284 | NULL | NULL | n.s. | n.s. | 199 | 199 | n.s. | n.s. | 332 | 332 | 210 | 210 |
| | | | | | | | | | | Scored: 181, 187 | | | | | | | | | | | | | | | | | | | | | | | | | | | | | | | | | | | |
| **PstS13 hybridization: Nucleus haplotype B (PstS7[1]/PstS10) and 2nd nucleus haplotype from PstS4** | | | | | | | | | | | | | | | | | | | | | | | | | | | | | | | | | | | | | | | | | | | | | |
| PstS13 genotype | Scored (MLG#21) | 224 | 224 | 172 | 176 | 196 | 196 | 150 | 150 | 169 | NULL | NULL | 336 | n.s. | n.s. | 257 | 255 | 228 | 226 | 315 | 315 | 304 | 307 | 332 | 332 | **337** | **331** | 287 | 287 | 176 | 170 | 272 | 284 | 230 | 230 | n.s. | n.s. | 205 | 199 | n.s. | n.s. | 223 | 232 | 210 | 210 |
| Haplotype B: PstS7/PstS10 | Genome blast & SSR assignment | | 224 | | 176 | | 196 | | 150 | | NULL | | 336 | | n.s. | | 255 | | 226 | | 315 | | 307 | | 332 | | **331** | | 287 | | 170 | | 284 | | NULL | | n.s. | | 199 | | n.s. | | 332 | | 210 |
| Haplotype A: Consistent with PstS4 | Inferred | 224 | | 172* | | 196 | | 150 | | 169 | | NULL | | n.s. | | 257 | | 228 | | 315 | | 304 | | 332 | | **337** | | 287 | | 176 | | 272* | | 230 | | n.s. | | 205 | | n.s. | | 223 | | 210 | |
| | | | | *uniquely shared by PstS13/PstS4 isolates | | | | | | | | | | | | | | | | | | | | | | | | | | | | *uniquely shared by PstS13/PstS4 isolates | | | | | | | | | | | | | |
| **PstS14 hybridization: Nucleus haplotype B (PstS7[1]/PstS10/ PstS13) and 2nd nuclear haplotype from PstS1_2** | | | | | | | | | | | | | | | | | | | | | | | | | | | | | | | | | | | | | | | | | | | | | |
| PstS14 genotype | Scored (MLG#138) | 227 | 224 | 174 | 176 | 196 | 196 | 150 | 150 | 171 | NULL | NULL | 336 | n.s. | n.s. | 257 | 255 | 228 | 226 | 318 | 315 | 304 | 307 | 332 | 332 | **337** | **331** | 287 | 287 | 179 | 170 | 293 | 284 | 230 | 230 | n.s. | n.s. | 202 | 199 | | n.s. | 323 | 332 | 210 | 210 |
| Haplotype B: PstS7/PstS10 | Genome blast & SSR assignment | | 224 | | 176 | | 196 | | 150 | | NULL | | 336 | | n.s. | | 255 | | 226 | | 315 | | 307 | | 332 | | **331** | | 287 | | 170 | | 284 | | NULL | | n.s. | | 199 | n.s. | n.s. | | 332 | | 210 |
| Haplotype A: PstS1_2 | Inferred and genome validated | 227 | | 174 | | 196 | | 150 | | 171 | | NULL | | n.s. | | 257 | | 228 | | 318 | | 304 | | 332 | | **337** | | 287 | | 179* | | 293 | | 230 | | n.s. | | 202 | | | n.s. | 323 | | 210 | |
| | | | | | | | | | | | | | | | | | | | | | | | | | | | | *uniquely shared by PstS14/PstS1_2 isolates | | | | | | | | | | | | | | | |

[1] PstS7 most likely parent based on epidemiological data

| | | | | | | | | | | | | | | | | | | | | | | | | | | | | | | | | | | | | | | | | | | | | | |
|---|---|---|---|---|---|---|---|---|---|---|---|---|---|---|---|---|---|---|---|---|---|---|---|---|---|---|---|---|---|---|---|---|---|---|---|---|---|---|---|---|---|---|---|---|---|
| **PstS18 hybridization: Nucleus haplotype A (PstS1_2) and 2nd nucleus haplotype from US isolate (MLG # 907) significantly different from named clonal groups (PstS0 - PstS18)** | | | | | | | | | | | | | | | | | | | | | | | | | | | | | | | | | | | | | | | | | | | | | |
| PstS18 genotype | Scored (MLG#906) | 227 | 224 | 174 | 178 | 196 | 196 | 150 | 147 | 171 | NULL | NULL | 336 | n.s. | n.s. | 257 | 255 | 228 | 226 | 318 | 318 | 304 | 307 | 332 | 332 | **337** | **357** | 287 | 284 | 179 | 170 | 293 | 284 | 230 | NULL | n.s. | n.s. | 202 | 199 | | n.s. | 323 | 332 | 210 | 210 |
| Haplotype A: PstS1_2 | inferred and genome validated | 227 | | 174 | | 196 | | 150 | | 171 | | NULL | | n.s. | | 257 | | 228 | | 318 | | 304 | | 332 | | **337** | | 287 | | 179 | | 293 | | 230 | | n.s. | | 202 | | | n.s. | 323 | | 210 | |
| Haplotype B: Other (MLG #907) | inferred | | 224 | | 178 | | 196 | | 147 | | NULL | | 336 | | | | 255 | | 226 | | 318 | | 307 | | 332 | | **357** | | 284 | | 170 | | 284 | | NULL | | | | 199 | | | | 332 | | 210 |

**Table S6** Overview of 419 people contributing to world-wide sampling of yellow rust between 2009 and 2023. When available, affiliations at time of sampling have been provided

| | Country | Collector | Institution |
|---|---|---|---|
| 1 | Afghanistan | Abdul Bari | |
| 2 | Afghanistan | Abdul Bari Stanikzai | |
| 3 | Afghanistan | Abdul Latif Rasekh | |
| 4 | Afghanistan | Abdul Raqeeb | |
| 5 | Afghanistan | Abdul Raqib Lodin | |
| 6 | Afghanistan | Ahmad Samin Samimy | |
| 7 | Afghanistan | Ahmadi Nabizada | |
| 8 | Afghanistan | Ahmadjan Noori | |
| 9 | Afghanistan | Baryalai Malekzada | |
| 10 | Afghanistan | Elias Mohmand | |
| 11 | Afghanistan | Gheyasuddin Ghanizada | |
| 12 | Afghanistan | Mahmoodian | |
| 13 | Afghanistan | Mirwais Amirzai | |
| 14 | Afghanistan | Mohaqiq | |
| 15 | Afghanistan | Noorul Haq | |
| 16 | Afghanistan | Raqib Lodin | |
| 17 | Afghanistan | Ravij Sharma | CIMMYT |
| 18 | Afghanistan | Saifudin Safi | |
| 19 | Afghanistan | Shakib Ataee | |
| 20 | Afghanistan | Shamsullhaq | |
| 21 | Afghanistan | Zemaray Ahmazada | |
| 22 | Algeria | Chaneze | |
| 23 | Argentina | Adelina Larsen | |
| 24 | Argentina | Agustin Bilbao | |
| 25 | Argentina | Agustín Pulido | MyCAI |
| 26 | Argentina | Alejandro Porfiri | |
| 27 | Argentina | Ana Rodriguez | |
| 28 | Argentina | Ana Storm | |
| 29 | Argentina | Andrea Rosso | Ceanagro, SA. |
| 30 | Argentina | Buck Semillas | |
| 31 | Argentina | Carina Cáceres | |
| 32 | Argentina | Carlos Grosso | VMV Siembras |
| 33 | Argentina | Carmona, Rancagua | |
| 34 | Argentina | Claudio Bosco | |
| 35 | Argentina | Cristian Babrilla | |
| 36 | Argentina | Cristina Palacios | |
| 37 | Argentina | Daniel Ploper | |
| 38 | Argentina | Diego Alvarez | FAUBA |
| 39 | Argentina | Duarte | DZD Agro |
| 40 | Argentina | Enrique Alberione | |
| 41 | Argentina | Fabricio Mock | BASF |
| 42 | Argentina | Franco Petrelli | |
| 43 | Argentina | Gustavo Duarte | El Ganado SRL |
| 44 | Argentina | Ignacio Erreguerena | |
| 45 | Argentina | Jonathan Damiani | Independent Consultant |
| 46 | Argentina | Julián Garcia, Oro Verde | Oro Verde |
| 47 | Argentina | Liliana Wehrhahne | |
| 48 | Argentina | Lucrecia Couretot | INTA Pergamino |
| 49 | Argentina | Magliano F | |
| 50 | Argentina | Manuela Gordo | |
| 51 | Argentina | Marcos Mitelsky | LIM Agro |
| 52 | Argentina | Margarita Sillon | |
| 53 | Argentina | Mariano Vence | CREA Loberías Grandes |
| 54 | Argentina | Mauro Montarini | |
| 55 | Argentina | Norma Formento | INTA Paraná |
| 56 | Argentina | Pablo Campos | |
| 57 | Argentina | Roxana Maumary | UNL |
| 58 | Argentina | Victoria Gonzales | |
| 59 | Azerbaijan | Beyhan Akin | CIMMYT, Turkiye |
| 60 | Azerbaijan | Konul Aslanova | Azerbaijan Research Institute of Crop Husbandry, Baku, Azerbaijan |
| 61 | Bhutan | Gordan Cisar | Cornell University |

| | | | |
|---|---|---|---|
| 62 | Bhutan | Namgay Om | NPPC |
| 63 | Bhutan | Sangay Tshewang | ARDC, Bajo |
| 64 | Bhutan | Sonam Dorji, Legjay | NPPC / ARDC Bajo |
| 65 | Bhutan | Tshomo | NPPC |
| 66 | Bhutan | Yeshey Dema | ARDC, Bajo |
| 67 | Brazil | Amarilis Barcellos | |
| 68 | Brazil | Ané Rosa | |
| 69 | Brazil | Marcia Chavez | |
| 70 | Brazil | Ricardo Castro | |
| 71 | Canada | Dean Landfood | |
| 72 | Chile | Ricardo Maiaga | |
| 73 | Chile | C. Jobet | |
| 74 | Chile | Carola Vera | |
| 75 | Chile | Erik Von Baer | |
| 76 | Chile | R. Galdames | |
| 77 | Chile | Ricardo Madariaga | |
| 78 | Denmark | Anders Almskov | Aarhus University |
| 79 | Denmark | Anders Kjær | Fjorland |
| 80 | Denmark | Anders Tobiasen | VKST |
| 81 | Denmark | Anne Ladegaard | Lemvigegnens Landboforening |
| 82 | Denmark | Annette V. Vestergård | SEGES |
| 83 | Denmark | Bent H. Hedegaard | Velas |
| 84 | Denmark | Camilla Beck Nielsen | Agrovi |
| 85 | Denmark | Chris K. Sorensen | Aarhus University |
| 86 | Denmark | Christian Møller Holm | Bornholms Landbrug & Fødevarer |
| 87 | Denmark | Christian Thormann Nielsen | ØkologiRådgivning Danmark |
| 88 | Denmark | Erik Silkjær Pedersen | Djursland Landboforening |
| 89 | Denmark | Finn Borum | Sejet Plant Breeding |
| 90 | Denmark | Ghita Nielsen | SEGES |
| 91 | Denmark | Gitte Skovgaard | Velas |
| 92 | Denmark | Hans Christian Jacobsen | VKST |
| 93 | Denmark | Hans Erik Larsen | Agrogården |
| 94 | Denmark | Henning Skov Andersen | |
| 95 | Denmark | Henrik Østergaard Nielsen | |
| 96 | Denmark | Irene Rasmussen | VKST |
| 97 | Denmark | Jan Lund | LandboNord |
| 98 | Denmark | Jens Kristian Steensen | Agrovi |
| 99 | Denmark | Jeppe Reitan | Nordic Seed |
| 100 | Denmark | Julie Torp-Thomsen | Østdansk Landboforening |
| 101 | Denmark | Kenneth Svensson | VKST |
| 102 | Denmark | Kirsten Larsen | Velas |
| 103 | Denmark | Kirsten S. Pedersen | LandboNord |
| 104 | Denmark | Kjeld Forsom | ØkologiRådgivning Danmark |
| 105 | Denmark | Lars B. Skovborg | LandboThy |
| 106 | Denmark | Lars Egelund Olsen | SEGES |
| 107 | Denmark | Lisa Munk | Copenhagen University |
| 108 | Denmark | Lise Nistrup Jørgensen | Aarhus University |
| 109 | Denmark | Lotte Olsen | Nordic Seed |
| 110 | Denmark | Mads Brandt | Landbrugsrådgivning Syd |
| 111 | Denmark | Margit Bæk Jensen | HortiAdvice A/S |
| 112 | Denmark | Martin Nielsen | |
| 113 | Denmark | Martin Ugilt Thomsen | Private |
| 114 | Denmark | Mehran Patpour | Aarhus University |
| 115 | Denmark | Michael Fleng | Sagro |
| 116 | Denmark | Michael Riis | Landbothy |
| 117 | Denmark | Michael Wang Lønbæk | Centrovice |
| 118 | Denmark | Ole Harild | Bornholms Landbrug & Fødevarer |
| 119 | Denmark | Peter Sivertsen | |
| 120 | Denmark | Pia Bay | SEGES |
| 121 | Denmark | Poul Christensen | Privat Planteavls Rådgivning |
| 122 | Denmark | Rasmus Hjortshøj | Sejet Plant Breeding |
| 123 | Denmark | Rikke B. heinfelt | VKST |
| 124 | Denmark | Susanne Sindberg | Tystofte Foundation |
| 125 | Denmark | Søren Banke | Sejet Plant Breeding |

| | | | |
|---|---|---|---|
| 126 | Denmark | Thomas Olsen | |
| 127 | Denmark | Thomas Wohlleben | Velas |
| 128 | Denmark | Torben Kejser | VKST |
| 129 | Egypt | Atef Shahin | Department of Wheat Diseases ResearchCairo, Egypt |
| 130 | Egypt | Doaa Ragheb El-Naggar | Plant Pathology Research Institute, Agricultural Research Centre, Giza, Egypt |
| 131 | Egypt | Essam Abdelhamid | |
| 132 | Egypt | Mamdouh Asmawi | |
| 133 | Egypt | Reda Ibrahim Nada Omara | Plant Pathology Research Institute, Agricultural Research Centre, Giza, Egypt |
| 134 | Egypt | Walid El.Orabey | Plant Pathology Research Institute, Agricultural Research Centre, Giza, Egypt |
| 135 | Eritrea | Ashmelash Wolday | Ministry of Agriculture, National Agricultural Research Institute (NARI) |
| 136 | Ethiopia | Almaz Weytso | EIAR, Ambo ARC |
| 137 | Ethiopia | Anduamlak A | EIAR, DebreMarkos ARC |
| 138 | Ethiopia | Asela Kesho | EIAR, Holeta ARC |
| 139 | Ethiopia | Ashenafi Gemechu | EIAR, Bishoftu ARC |
| 140 | Ethiopia | Ayele Badebo | CIMMYT |
| 141 | Ethiopia | Bedada Girma | EIAR, Kulumsa ARC |
| 142 | Ethiopia | Bekele Abeyo | CIMMYT |
| 143 | Ethiopia | Bekele Hundie | EIAR, Kulumsa ARC |
| 144 | Ethiopia | Belachew Bekele | EIAR, Debremarkos ARC |
| 145 | Ethiopia | Dagne Abu | OARI, Bore ARC |
| 146 | Ethiopia | Dawit Asnake | EIAR Kulumsa ARC |
| 147 | Ethiopia | Dereje Amare | Ambo ARC |
| 148 | Ethiopia | Fikrte Yirga | EIAR, Kulumsa ARC |
| 149 | Ethiopia | Gabisa G. | EIAR, Jimma ARC |
| 150 | Ethiopia | Gebre | EIAR, Debre Zeit ARC |
| 151 | Ethiopia | Gerald Blasch | CIMMYT |
| 152 | Ethiopia | Getahun Tena | EIAR, Holeta ARC |
| 153 | Ethiopia | Getaneh Woldeab | EIAR, Ambo ARC |
| 154 | Ethiopia | Getenesh Demsie | EIAR, Kulumsa ARC |
| 155 | Ethiopia | Getnet Muche | EIAR, Kulumsa ARC |
| 156 | Ethiopia | Girma Abebe | EIAR, Holeta ARC |
| 157 | Ethiopia | Girma Teshome | OARI, Bore ARC |
| 158 | Ethiopia | Habtamu Tesfaye | EIAR, Bishoftu ARC |
| 159 | Ethiopia | Hailu Negassa | EIAR, Jimma ARC |
| 160 | Ethiopia | Hawila Tefaye | EIAR, Kulumsa ARC |
| 161 | Ethiopia | Jemal Tola | EIAR, Ambo ARC |
| 162 | Ethiopia | Kabna Asefa | OARI, Bore ARC |
| 163 | Ethiopia | Girma Teshome | EIAR, Holeta |
| 164 | Ethiopia | Mitiku Kebede | SARI, Werabe ARC |
| 165 | Ethiopia | Kitessa Gutu | EIAR, Ambo ARC |
| 166 | Ethiopia | Lidiya Tesfaye | EIAR, Kulumsa ARC |
| 167 | Ethiopia | Luche Soboka | EIAR, Ambo ARC |
| 168 | Ethiopia | Mathios Ashamo | SARI, Hawassa ARC |
| 169 | Ethiopia | Mequanint Andualem | ARARI, Adet ARC |
| 170 | Ethiopia | Misgana Mitiku | SARI, Werabe ARC |
| 171 | Ethiopia | Netsanet Bacha | EIAR |
| 172 | Ethiopia | Nurhussein Seid | EIAR, Werer ARC |
| 173 | Ethiopia | Solomon Shibeshi | SARI, Hawassa ARC |
| 174 | Ethiopia | Tamene Mideksa | EIAR, Holeta |
| 175 | Ethiopia | Tamirat Negash | EIAR Kulumsa ARC |
| 176 | Ethiopia | Tekilu | EIAR, Ambo ARC |
| 177 | Ethiopia | Tilahun Bayisa | OARI, Sinana ARC |
| 178 | Ethiopia | Tizazu | EIAR, Ambo ARC |
| 179 | Ethiopia | TsegaAb Tesfaye | EIAR, Ambo ARC |
| 180 | Ethiopia | Yared Tesfaye | SARI, Bore ARC |
| 181 | Ethiopia | Yitagesu Tadesse | EIAR, Holeta ARC |
| 182 | Ethiopia | Yoseph Alemayehu | CIMMYT |
| 183 | Ethiopia | Zenebe Hailu | EIAR |
| 184 | Ethiopia | Zenebe Wubisher | EIAR, Jimma ARC |
| 185 | Ethiopia | Zerihun Eshetu | OARI Sinana ARC |
| 186 | Ethiopia | Zerihun Tadesse | EIAR, Kulumsa ARC |
| 187 | Finland | Auli Kedonperä | Luke |
| 188 | Finland | Marja Jalli | Luke |
| 189 | Iran | Farzad Afshari | Seed and Plant Improvement Institute (SPII), Karaj, Iran |

| | | | |
|---|---|---|---|
| 190 | Iran | Goodarz Najafian | Seed and Plant Improvement Institute (SPII), Karaj, Iran |
| 191 | Iran | Mahboobeh Yazdani | |
| 192 | Iraq | Abid Al.Hameed Fayadh | |
| 193 | Iraq | Ahmed Neama Jwad | |
| 194 | Iraq | Dheyaa Mohsin Ali | |
| 195 | Iraq | Dhia Muhsen Ali | |
| 196 | Iraq | Emad Al-Maaroof | Sulaimani College of Agricultural Engineering Sciences, University of Sulaimani, Iraq |
| 197 | Iraq | Hatem Mahmood Hassan | |
| 198 | Iraq | Hatim Hussien | |
| 199 | Iraq | Hazha Abdul Karim | |
| 200 | Iraq | Laith Husain | |
| 201 | Iraq | Nabaz Rashid | |
| 202 | Iraq | Sarkawt Salih | University of Sulaimani |
| 203 | Kazakhstan | Alma Kohkmetova | |
| 204 | Kazakhstan | Gulzat Yessenbekova | |
| 205 | Kenya | Godwin Macharia | KALRO, Njoro |
| 206 | Kenya | R. Wanyera | KALRO, Njoro |
| 207 | Kenya | S. Kilonzo | KALRO, Njoro |
| 208 | Kenya | Zak Pretorius | UFS, South Africa |
| 209 | Kenya | Zennah Kosgey | KALRO, Njoro |
| 210 | Lebanon | Rola ElAmil | |
| 211 | Mexico | Neela Quershi | CIMMYT |
| 212 | Mexico | Pablo Espejel | CIMMYT |
| 213 | Mexico | Sridhar Bhavani | CIMMYT |
| 214 | Morocco | Adbdelhamid Ramdani | Institut Nationale de la Recherche Agronomique (INRA-Meknes) |
| 215 | Morocco | Ezzahiri Brahim | Hassan II Agronomic and Veterinary Institute, Rabat, Morocco |
| 216 | Morocco | Ilyass Maaf | ICARDA, Morocco |
| 217 | Morocco | Muamar Al-jaboobi | ICARDA, Morocco |
| 218 | Morocco | Seid Kemal | ICARDA, Morocco |
| 219 | Nepal | Basistha Acharya | NPPRC, NARC |
| 220 | Nepal | Bhajuman Maharjan | |
| 221 | Nepal | Bhajuman, Bhojraj | |
| 222 | Nepal | Dhurba Thapa | NPBGC, NARC |
| 223 | Nepal | EK Bra. Shrestha | NARC |
| 224 | Nepal | Khem Pant | NWP, Bhairahawa, NARC |
| 225 | Nepal | Laxman Aryal | NWP, Bhairahawa, NARC |
| 226 | Nepal | Laxmi Shrestha | |
| 227 | Nepal | Mahesh Subedi | NPBGC, NARC |
| 228 | Nepal | Pavitri Joshi | |
| 229 | Nepal | Prem Bahadur Magar | NPPRC, NARC |
| 230 | Nepal | Roshan Basnet | NWP, Bhairahawa, NARC |
| 231 | Nepal | Sunita Adhikari | |
| 232 | Nepal | Suraj Baidya | NPPRC, NARC |
| 233 | Norway | Andrea Ficke | NIBIO |
| 234 | Norway | B. Lesteberg | Farmer |
| 235 | Norway | Chloe Grieu | NIBIO |
| 236 | Norway | H. Mørk | Farmer |
| 237 | Norway | H. Thirud | Farmer |
| 238 | Norway | Inga Holt | NLR |
| 239 | Norway | Ingvild Evju | NLR |
| 240 | Norway | Jafar Razaghian | NIBIO |
| 241 | Norway | K. Eskerud | Farmer |
| 242 | Norway | L. O. Rokstad | Farmer |
| 243 | Norway | Margit Kim | Graminor |
| 244 | Norway | Min Lin | NMBU |
| 245 | Norway | Morten Lillemo | NMBU |
| 246 | Norway | Nils Kristian Aker | NLR |
| 247 | Norway | O.K. Fevik | Farmer |
| 248 | Norway | Ole Jakob Ulberg | NLR |
| 249 | Norway | Rune Karlsen | NLR |
| 250 | Norway | Sanna Persson | NLR |
| 251 | Norway | Silja Valand | NLR |
| 252 | Norway | Thomas Julseth Brown | NLR |
| 253 | Norway | Tina Fallet | Strand Unikorn |

| | | | |
|---|---|---|---|
| 254 | Norway | Unni Abrahamsen | NIBIO |
| 255 | Norway | Unni Roed | NLR |
| 256 | Norway | Ø. Jensen | Farmer |
| 257 | Pakistan | Munsif | University of Agriculture, Peshawar 25130, Khyber Pakhtunkhwa |
| 258 | Pakistan | Noor | University of Agriculture, Peshawar 25130, Khyber Pakhtunkhwa |
| 259 | Pakistan | Rameez Kahn | University of Agriculture, Peshawar 25130, Khyber Pakhtunkhwa |
| 260 | Pakistan | Muhammad Fayyaz | Crop Diseases Research Institute, NARC, Islamabad |
| 261 | Pakistan | Sajid Ali | University of Agriculture, Peshawar 25130, Khyber Pakhtunkhwa |
| 262 | Paraguay | Marta Fernandez | IPTA |
| 263 | Rwanda | Aloys | |
| 264 | Rwanda | Annualite | |
| 265 | Rwanda | Athanase | |
| 266 | Rwanda | Eliane | |
| 267 | Rwanda | Felicien | |
| 268 | Rwanda | Innocent Habarurema | |
| 269 | Rwanda | Jean Marie | |
| 270 | Rwanda | Obed | |
| 271 | Rwanda | Theoneste | |
| 272 | Sweden | Agnes Jonsson | Jordbruksverket |
| 273 | Sweden | Agnes Radovic | Jordbruksverket |
| 274 | Sweden | Alexia von Ehrenheim | Jordbruksverket |
| 275 | Sweden | Alf Djurberg | Jordbruksverket |
| 276 | Sweden | Amanda Ahlqvist | Jordbruksverket |
| 277 | Sweden | Anders Arvidsson | Jordbruksverket |
| 278 | Sweden | Anders Karlsson | Jordbruksverket |
| 279 | Sweden | Anders Lindgren | Jordbruksverket |
| 280 | Sweden | Anna Berlin | Jordbruksverket |
| 281 | Sweden | Anna Gerdtsson | Jordbruksverket |
| 282 | Sweden | Anna Magnusson | Jordbruksverket |
| 283 | Sweden | Anna Olsson | Jordbruksverket |
| 284 | Sweden | Anna Pers | Jordbruksverket |
| 285 | Sweden | Anna von Heideken | Jordbruksverket |
| 286 | Sweden | Anna-Karin Krijger | Jordbruksverket |
| 287 | Sweden | Annika Sohlman | Jordbruksverket |
| 288 | Sweden | Anton Hampl | Jordbruksverket |
| 289 | Sweden | Aron Westlin | Jordbruksverket |
| 290 | Sweden | Camilla Broms | Jordbruksverket |
| 291 | Sweden | Caroline Jøngren | Jordbruksverket |
| 292 | Sweden | Cecilia Lerenius | Jordbruksverket |
| 293 | Sweden | Cecilia Söderlind | Jordbruksverket |
| 294 | Sweden | Charlotte Norén | Jordbruksverket |
| 295 | Sweden | Clara Andersson | Jorbruksverket |
| 296 | Sweden | Danira Behaderovic | Jordbruksverket |
| 297 | Sweden | Ebba Hellstrand | Jordbruksverket |
| 298 | Sweden | Elin Almén | Jordbruksverket |
| 299 | Sweden | Elin Nilsson | Jordbruksverket |
| 300 | Sweden | Elin Strömberg | Jorbruksverket |
| 301 | Sweden | Elisabeth Bölenius | Jordbruksverket |
| 302 | Sweden | Elisabeth Lövstad | Jordbruksverket |
| 303 | Sweden | Emil | Jordbruksverket |
| 304 | Sweden | Emmy Johansson | Jordbruksverket |
| 305 | Sweden | Erling Christensson | Jordbruksverket |
| 306 | Sweden | Eva Mellqvist | Jordbruksverket |
| 307 | Sweden | Fia Birch-Jensen | Jordbruksverket |
| 308 | Sweden | Frans Johnson | Jordbruksverket |
| 309 | Sweden | Frida Erlöv | Jordbruksverket |
| 310 | Sweden | Gunilla Berg | Jordbruksverket |
| 311 | Sweden | Gunnel Andersson | Jordbruksverket |
| 312 | Sweden | Göran Gustafsson | Jordbruksverket |
| 313 | Sweden | Hanna Johansson | Jordbruksverket |
| 314 | Sweden | Hans Hedström | Jordbruksverket |
| 315 | Sweden | Jaokim Hermansson | Jordbruksverket |
| 316 | Sweden | Jenny Knutsson | Jordbruksverket |
| 317 | Sweden | Johan Andersson | Jordbruksverket |

| | | | |
|---|---|---|---|
| 318 | Sweden | Johanna Holmblad | Jordbruksverket |
| 319 | Sweden | Johanna Lindgren | Jordbruksverket |
| 320 | Sweden | Jonas Törngren | Jordbruksverket |
| 321 | Sweden | Jonathan Sjöström | Jordbruksverket |
| 322 | Sweden | Julia Dahlqvist | Jordbruksverket |
| 323 | Sweden | Karin Andersson | Jordbruksverket |
| 324 | Sweden | Kristian Barck | Jordbruksverket |
| 325 | Sweden | Kristian Jochnick | Jordbruksverket |
| 326 | Sweden | Lars Johansson | Jordbruksverket |
| 327 | Sweden | Lina Norrlund | Jordbruksverket |
| 328 | Sweden | Linda Geijersstam | |
| 329 | Sweden | Linnea Aronsson | Jordbruksverket |
| 330 | Sweden | Lisbeth Bergh | Jordbruksverket |
| 331 | Sweden | Louisa Bergkvist | Jordbruksverket |
| 332 | Sweden | Louise Adler | Jordbruksverket |
| 333 | Sweden | Louise Aldén | Jordbruksverket |
| 334 | Sweden | Lovisa Eriksson | Jordbruksverket |
| 335 | Sweden | Lukas Hallberg | Jordbruksverket |
| 336 | Sweden | M. Thorngren | Jordbruksverket |
| 337 | Sweden | Mahboobeh | SLU |
| 338 | Sweden | Mahbubjon Ramatov | SLU |
| 339 | Sweden | Mariann Wikström | Jordbruksverket |
| 340 | Sweden | Marie Björs | Jordbruksverket |
| 341 | Sweden | Markus Eriksson | Västskyddscentralen |
| 342 | Sweden | Mats Selin | Jordbruksverket |
| 343 | Sweden | Mikael Nilsson | LAAP |
| 344 | Sweden | Nils Wiklund | Jordbruksverket |
| 345 | Sweden | Nils, Johan | Jordbruksverket |
| 346 | Sweden | Oscar Andersson | Jordbruksverket |
| 347 | Sweden | Oskar Björing | Jordbruksverket |
| 348 | Sweden | Oskar Gustafsson | Jordbruksverket |
| 349 | Sweden | Petter Gustafsson | Jordbruksverket |
| 350 | Sweden | Rebecka Östlund | Jordbruksverket |
| 351 | Sweden | Robert Dinwiddie | Jordbruksverket |
| 352 | Sweden | Rolf Lindholm | Jorbruksverket |
| 353 | Sweden | Sara Furenhed | Jordbruksverket |
| 354 | Sweden | Sara Wallemyr | Jordbruksverket |
| 355 | Sweden | Sofie Pålsson | Jordbruksverket |
| 356 | Sweden | Susanna Waara | Jordbruksverket |
| 357 | Sweden | Therese Christerson | Jordbruksverket |
| 358 | Sweden | Lina Norrlund | Jordbruksverket |
| 359 | Syria | G. Mhairy | Plant Protection Directorate, Ministry of Agriculture and Agrarian Reform, Damascus, Syria |
| 360 | Syria | K. Heimoun | Plant Protection Directorate, Ministry of Agriculture and Agrarian Reform, Damascus, Syria |
| 361 | Syria | Maha Al Ahmed | ICARDA |
| 362 | Syria | Majd Jamal | ICARDA-Syria |
| 363 | Syria | Meni | |
| 364 | Syria | S.A. Ghorrah | Plant Protection Directorate, Ministry of Agriculture and Agrarian Reform, Damascus, Syria |
| 365 | Tajikistan | A. Morgounov | CIMMYT |
| 366 | Tajikistan | Eshonova | |
| 367 | Tajikistan | Huseynov | |
| 368 | Tajikistan | Mahbubjon Ramatov | SLU |
| 369 | Tanzania | Charles Msuya | TARI |
| 370 | Tanzania | Christina Petro Buluba | TARI, Selian |
| 371 | Tanzania | Hosea Nyato | TARI, Uyole |
| 372 | Tanzania | Michael Paul | Davinco |
| 373 | Tanzania | Moses Mbwana | TARI |
| 374 | Tanzania | Okinyi, Moses | TARI |
| 375 | Tanzania | Rose Mongi, Nelson | TARI, Uyole |
| 376 | Tanzania | Salome W. Munissi | TARI, Selian |
| 377 | Tanzania | Seni Marco | TARI, Selian |
| 378 | Tunisia | Amor Yahyaoui | ICARDA |
| 379 | Tunisia | Mohamed Salah Gharbi | |
| 380 | Tunisia | Sarrah Ben M'Barek | Regional Field Crops research Center of Beja |
| 381 | Tunisia | Tarek Jarrahi | Institut National des Grandes Cultures (INGC), Tunisia |

| | | | |
|---|---|---|---|
| 382 | Tunisia | Wided Abdedeyam | Regional Field Crops research Center of Beja |
| 383 | Türkiye | Ali Kadiroğlu | Regional Cereal Rust Research Center, Aegean Agricultural Research Institute, Izmir, Türkiye |
| 384 | Türkiye | Beyhan Akin | CIMMYT, Turkiye |
| 385 | Türkiye | Ceren Cer | Directorate Plant Protection Research Institute, Erzene, P.O. Box 35040, Bornova, Izmir, Turkey |
| 386 | Türkiye | Emine Burcu | Field Crops Central Research Institute, Ankara, Türkiye |
| 387 | Türkiye | Ezgi Kurtulus | Syngenta Turkey, Crop Protection Department, Konak-Izmir, 35170, Türkiye |
| 388 | Türkiye | Gürkan Basbagci | Bati Akdeniz Agricultural Research Institute, Department of Plant Health, Antalya, Türkiye |
| 389 | Türkiye | Hakan Hekiman | Regional Cereal Rust Research Center, Aegean Agricultural Research Institute, Izmir, Türkiye |
| 390 | Türkiye | Hakan Hekiman | Regional Cereal Rust Research Center, Aegean Agricultural Research Institute, Izmir, Türkiye |
| 391 | Türkiye | Handan Kavaz | Viticulture Research Institute, Plant Protection Department Manisa, Türkiye |
| 392 | Türkiye | Hatice Geren | Aegean Agricultural Research Institute, P.K. 9, Menemen, Izmir, Türkiye |
| 393 | Türkiye | Izzet Ozseven | Regional Cereal Rust Research Center, Aegean Agricultural Research Institute, Izmir, Türkiye |
| 394 | Türkiye | Omer M. Ozturk | Directorate of Provincial Agriculture and Forestry, Pamukkale P.O. Box 20150, Denizli, Turkey |
| 395 | Türkiye | Yesim Egerci | Directorate Plant Protection Research Institute, Erzene, P.O. Box 35040, Bornova, Izmir, Turkey |
| 396 | Turkey | Zafer Mert | Central Research Institute for Field Crops, Ankara, Turkey |
| 397 | Uganda | Arthur Wasukira | BugiZardi, NARO |
| 398 | Uganda | Bonny | BugiZardi, NARO |
| 399 | Uganda | Bosco Chemayek | BugiZardi, NARO |
| 400 | Uganda | Eric | BugiZardi, NARO |
| 401 | Uganda | Kenneth | BugiZardi, NARO |
| 402 | Uganda | Lawrence Owere | BugiZardi, NARO |
| 403 | Uganda | Wobibi | BugiZardi, NARO |
| 404 | Uganda | Woniala | BugiZardi, NARO |
| 405 | Uruguay | F. Pereira | INIA |
| 406 | Uruguay | R. García | INIA |
| 407 | Uruguay | Silvia German | INIA |
| 408 | USA | Gene Milus | University of Arkansas, USA |
| 409 | Uzbekistan | Baboev S | |
| 410 | Uzbekistan | Ram Chane Sharma | ICARDA, Tashkent, Uzbekistan |
| 411 | Uzbekistan | Safar Alikulov | |
| 412 | Uzbekistan | Zafar Ziyaev | Uzbek Scientific and Production Center of Agriculture, Tashkent, Uzbekistan |
| 413 | Yemen | Aref Alshamiri | |
| 414 | Yemen | Mohamed Alsadi | |
| 415 | Yemen | Musaed Eisa | |
| 416 | Yemen | Rashad Basha | |
| 417 | Yemen | Sailan | |
| 418 | Yemen | Wajeeh Almutawakel | |
